# A *Bacillus anthracis* Genome Sequence from the Sverdlovsk 1979 Autopsy Specimens

**DOI:** 10.1101/069914

**Authors:** Jason W. Sahl, Talima Pearson, Richard Okinaka, James M. Schupp, John D. Gillece, Hannah Heaton, Dawn Birdsell, Crystal Hepp, Viacheslav Fofanov, Ramón Noseda, Antonio Fasanella, Alex Hoffmaster, David M. Wagner, Paul Keim

**Author notes:** Address correspondence to Paul Keim. J.S. and T.P. contributed equally to this report.

## Abstract

Anthrax is a zoonotic disease that occurs naturally in wild and domestic animals but has been used by both state-sponsored programs and terrorists as a biological weapon. The 2001 anthrax letter attacks involved less than gram quantities of *Bacillus anthracis* spores while the earlier Soviet weapons program produced tons. A Soviet industrial production facility in Sverdlovsk proved deficient in 1979 when a plume of spores was accidentally released and resulted in one of the largest known human anthrax outbreak. In order to understand this outbreak and others, we have generated a *B. anthracis* population genetic database based upon whole genome analysis to identify all SNPs across a reference genome. Only ~12,000 SNPs were identified in this low diversity species and represents the breadth of its known global diversity. Phylogenetic analysis has defined three major clades (A, B and C) with B and C being relatively rare compared to A. The A clade has numerous subclades including a major polytomy named the Trans-Eurasian (TEA) group. The TEA radiation is a dominant evolutionary feature of *B. anthracis*, many contemporary populations, and must have resulted from large-scale dispersal of spores from a single source. Two autopsy specimens from the Sverdlovsk outbreak were deeply sequenced to produce draft *B. anthracis* genomes. This allowed the phylogenetic placement of the Sverdlovsk strain into a clade with two Asian live vaccine strains, including the Russian Tsiankovskii strain. The genome was examined for evidence of drug resistance manipulation or other genetic engineering, but none was found. Only 13 SNPs differentiated the virulent Sverdlovsk strain from its common ancestor with two vaccine strains. The Soviet Sverdlovsk strain genome is consistent with a wild type strain from Russia that had no evidence of genetic manipulation during its industrial production. This work provides insights into the world's largest biological weapons program and provides an extensive *B. anthracis* phylogenetic reference valuable for future anthrax investigations.

**Importance:** The 1979 Russian anthrax outbreak resulted from an industrial accident at the Soviet anthrax spore production facility in the city of Sverdlovsk. Deep genomic sequencing of two autopsy specimens generated a draft genome and phylogenetic placement of the Soviet Sverdlovsk anthrax strain. While it is known that Soviet scientists had genetically manipulated *Bacillus anthracis*, with the potential to evade vaccine prophylaxis and antibiotic therapeutics, there was no genomic evidence of this from the Sverdlovsk production strain genome. The whole genome SNP genotype of the Sverdlovsk strain was used to precisely identify it and its close relatives in the context of an extensive global *B. anthracis* strain collection. This genomic identity can now be used for forensic tracking of this weapons material on a global scale and for future anthrax investigations.

## Introduction

Anthrax is a zoonotic disease caused by *Bacillus anthracis* with a relatively small impact on global human health, but it has become notorious and widely feared due to its use and potential as a biological weapon. In its spore form, the bacterium represents a highly stable quiescent entity that is capable of surviving for decades, a critical part of its ecology, global distribution, evolution and infectivity. The vegetative phase allows for cellular proliferation following spore germination in a host animal. The vegetative form expresses specific mechanisms for avoiding the innate host immunity with some of these encoded on two large virulence plasmids – pXO1 and pXO2 (Mock and Fouet 2001). Adaptive immunity can be highly effective at preventing disease and, interestingly, anthrax was the first bacterial disease mitigated with a vaccine (Tigertt 1980). Vaccine development for this pathogen is an important veterinary and public health measure, but research with a potential weapon of mass destruction (WMD) unfortunately can also lead to highly similar research supporting pathogen weaponization. Therefore, the treaty created by the Biological Weapons Convention of 1975 with 175 State Parties prohibited all offensive efforts with any biological agent, including anthrax (Affairs 2016).

The *B. anthracis* spore stability, potential for aerosolization, and its ability to cause acute pulmonary disease have historically led to multiple nations weaponizing this bacterium. It is well documented that large-scale production of spores was accomplished by the United States, the United Kingdom and the Soviet Union (Leitenberg, Zilinskas et al. 2012). Industrial spore production involves numerous quality control features to ensure spore stabilization, particle size, and the retention of virulence with extensive growth. These state sponsored programs were to cease with the Biological Weapons Convention of 1975. However, there are least two recent examples of anthrax spores being used in biological attacks: the Aum Shinrikyo cult attempted a liquid dispersal of *B. anthracis* in 1993 (Takahashi, Keim et al. 2004), and the 2001 US anthrax letters that killed five and sickened an additional 17 (Jernigan, Raghunathan et al. 2002).

The offensive anthrax weapons development programs were stopped in the US and UK in the 1960s, but continued covertly in the Soviet Union for at least another 20 years (Leitenberg, Zilinskas et al. 2012). Soviet, and later Russian, research on anthrax included projects to genetically modify *B. anthracis* strains. First, antibiotic resistance was genetically engineered into the vaccine strain STI-1 using recombinant DNA and a plasmid vector (Stepanov, Marinin et al. 1996). This effort resulted in multidrug resistance to penicillin, rifampicin, tetracycline, chloramphenicol, macrolides and lyncomycin with retention of normal colony morphology (Stepanov, Marinin et al. 1996). The stated goal of this research was the development of novel vaccines that allowed the simultaneous use of a live-vaccine strain and antibiotics in the case of human exposure. Without the drug resistant live-vaccine strain, long-term antibiotic therapy is required. Secondly, the program genetically engineered hemolytic properties from *B. cereus* into *B. anthracis* by the transfer of cereolysin AB genes into the STI-1 strain, again via a recombinant plasmid (Pomerantsev, Staritsin et al. 1997). This genetic change resulted in a strain with unique pathogenic features that could overcome the standard STI-1 vaccine protection in animal studies. The generation of a hemolytic *B. anthracis* strain was ostensibly for research purposes to understand basic host immunomodulation during anthrax, yet yielded a strain and strategy that could defeat vaccine protection. Manipulating the *B. anthracis* genome to change its phenotypic properties can and has been accomplished, raising concerns about dual use.

Evidence of the Soviet anthrax program’s continuation and its scale were revealed by the 1979 industrial accident in Sverdlovsk USSR (now known as Ekaterinburg) where at least 66 people died of inhalational anthrax (Meselson, Guillemin et al. 1994). This event has been shrouded in mystery with governmental denials and little public investigation, but it does represent one of the largest known human inhalational anthrax outbreak in history (Leitenberg, Zilinskas et al. 2012). According to local sources (Alibek and Handelman 1999, Leitenberg, Zilinskas et al. 2012), in early April 1979 safety air filters were compromised during routine maintenance at the Ministry of Defense’s (MOD) Scientific Research Institute of Microbiology (SRIM) spore production facility, known as Compound 19. This resulted in a plume of spores that spread downwind and caused human anthrax cases up to 4 km away and animal cases up to 50 km away (Meselson, Guillemin et al. 1994). Russian pathologists investigated these deaths and generated formalin-fixed tissues from multiple victims for analysis. These specimens showed evidence of anthrax (Abramova, Grinberg et al. 1993) and along with later PCR-based DNA analyses (Jackson, Hugh-Jones et al. 1998, Price, Hugh-Jones et al. 1999, Okinaka, Henrie et al. 2008) that detected *B. anthracis*, confirming that this cluster of deaths was indeed due to anthrax.

Here we have continued the Sverdlovsk anthrax investigation through deep sequencing of the formalin-fixed tissues from two of the victims to generate a draft genomic sequence of the infecting *B. anthracis* strain. In this paper, we also report the phylogenetic analysis of SNPs discovered among 193 whole genome sequences, which provided a phylogenetic context for analysis of the Sverdlovsk samples and can be used for similar analysis of other samples of interest. This provides a high-resolution analysis with detailed clade and subclade structures defined by a curated SNP database. SNP genotyping accurately places the Sverdlovsk strain into a subclade defined by the Tsiankovskii vaccine strain. We also examine the genome sequences for evidence of genetic engineering and adaptation to large production biology. The results demonstrate the power of combining modern molecular biology methods with a high-resolution curated SNP database in order to analyze a *B. anthracis* strain involved in a historic anthrax incident.

## Methods Section

### Sverdlovsk Specimen DNA Sequencing

DNA was extracted from paraffin embedded formalin-fixed tissues from two victims as previously described (Jackson, Hugh-Jones et al. 1998). These extracts were characterized by qPCR (Okinaka, Henrie et al. 2008) and the two samples (Svd-1: 7.RA93.15.15, spleen; Svd-2: 21.RA93.38.4, lymph node) with the lowest Ct values were subjected to Illumina sequencing, first on a MiSeq and later on a HiSeq 2000. Sequencing libraries were constructed using the standard Kapa Biosystems Illumina NGS Library reagent kit (cat# KK8232, Kapa Biosystems, Boston, MA), using 12 cycles in the final amplification reaction. Due to the highly degraded nature of the input DNA, fragment size selection prior to library preparation targeted fragments <500 bp. Both samples yielded libraries with enough material for sequencing, and were pooled and then sequenced using an entire MiSeq 600 cycle paired end run with V3 chemistry. This same pool was subsequently sequenced on a HiSeq 2000, using two lanes.

### Sequence analysis

Sequencing adapters were trimmed from reads with Trimmomatic (Bolger, Lohse et al. 2014). For SNP discovery, reads were aligned against the finished genome of the Ames Ancestor (NC_007530, NC_007322, NC_007323) with BWA-MEM (Li 2013) and SNPs were called with the UnifiedGenotyper method in GATK (McKenna, Hanna et al. 2010, DePristo, Banks et al. 2011). These methods were wrapped by the NASP pipeline (http://tgennorth.github.io/NASP/) (Sahl, Lemmer et al. 2016). Functional information was applied to SNPs with SnpEff (Kent 2002).

### Error profile analysis

To understand the error profiles in the Sverdlovsk genomes, reads were aligned against Ames Ancestor with BWA-MEM and for each position, the number of alleles that conflicted with the dominant allele were divided by the total number of bases at the position; this value was considered the per base error rate. As a control, this procedure was also performed for a genome (A0362) in the same phylogenetic group. Error rates were binned into different categories and represented as a histogram (Figure S1).

### Genome Assembly

To obtain a draft genome assembly, reads from both victims were combined and assembled with SPAdes v. 3.6.0 (Bankevich, Nurk et al. 2012). The first 200 bases of each contig were aligned against the GenBank (Benson, Karsch-Mizrachi et al. 2012) nt database with BLASTN (Altschul, Gish et al. 1990) to identify contigs not associated with *B. anthracis*; contigs that significantly aligned against human sequence were removed from the assembly. The contiguity of the assembly was then improved through a reference guided approach with AlignGraph (Bao, Jiang et al. 2014), using Ames Ancestor as the reference. The assembly was polished with Pilon v. 1.3.0 (Walker, Abeel et al. 2014), resulting in 128 contigs. A dotplot analysis using mummerplot (Delcher, Salzberg et al. 2003) was used to examine the synteny against Ames Ancestor as the reference.

### Phylogenetic Reconstructions

We compared the genomes of 193 strains of *B. anthracis* (Table S1) against Ames Ancestor to find SNPs (Table S2) using the In Silico Genotyper (Sahl, Beckstrom-Sternberg et al. 2015) and the Northern Arizona SNP Pipeline (Sahl, Lemmer et al. 2016). All SNP loci, even those that are missing in some of the genomes, were retained for phylogenetic analyses. We used parsimony criteria and a heuristic search with default options using PAUP 4.0b10 (Wilgenbusch and Swofford 2003) to infer phylogenetic trees. We report homoplasy using the consistency index as a measure of accuracy (Archie 1996) as bootstrapping is a poor measurement of accuracy for trees with little homoplasy (Felsenstein 1985) in clonal organisms (Pearson, Busch et al. 2004, Pearson, Okinaka et al. 2009). It should be noted however that the consistency index is influenced by the number of taxa impacting, direct comparisons across trees (Archie 1989). The phylogeny for all *B. anthracis* genomes was rooted according to Pearson et al. (Pearson, Busch et al. 2004). Trees of individual clades and subclades were rooted using a *B. anthracis* strain from another clade or the first strain to diverge from the rest of the group as determined by the overall phylogeny of *B. anthracis*. Phylogenetic branches were named according to precedent (Van Ert, Easterday et al. 2007) and designated on trees (Figures S2-S12). In short, each branch contains a prefix “A.Br”, “B.Br”, “A/B.Br”, or “C.Br”, depending on the major clade designation, followed by an assigned number based upon the order of branch discovery within each of the major clades. This method maintains the branch name from previous publications and allows for the identification of novel branches. However, branch numbers of adjacent branch numbers will often not be contiguous. For each SNP, the branches on which character state changes occurred, as determined by PAUP (Wilgenbusch and Swofford 2003) using the DescribeTrees command, is listed in the supplemental material (Table S3).

For evolutionarily stable characters such as SNPs found in clonal organisms like *B. anthracis*, a single locus can define a branch and thus serve as a “canonical SNP” (Keim, Van Ert et al. 2004, Pearson, Busch et al. 2004, Van Ert, Easterday et al. 2007, Pearson, Okinaka et al. 2009). As such, the character states of only a small number of SNP loci need to be interrogated in order to place an unknown strain into the established phylogenetic order. The list of SNPs on each branch (Table S3) thus serves as a resource of signatures that can be used to define a branch. However, new genome sequences will cause existing branches to be split, requiring additional branch names and updating the branch designation of these SNPs.

### Data accession

All reads were submitted to the NCBI Sequence Read Archive for 21.RA93.38.4 (SRR2968141, SRR2968216) and 7.RA93.15.15 (SRR2968143, SRR2968198). Data for all other genomes was deposited under accession SRP066845.

## Results

### A High Resolution Reference Phylogeny

We have constructed a high-resolution reference phylogeny from a large global *B. anthracis* strain collection. This is presented with collapsed clades (Fig. 1) to illustrate the overall phylogenetic structure but with complete branching details and annotated SNPs in the supplemental material (Figs. S2-S12). The global phylogeny is comprised of genomes from 193 strains (Table S1) that represent the global diversity as defined by other subtyping methods such as MLVA (Keim, Price et al. 2000) and canonical SNPs (Van Ert, Easterday et al. 2007, Marston, Allen et al. 2011, Price, Seymour et al. 2012, Khmaladze, Birdsell et al. 2014). Genomic sequence comparisons yielded 11,989 SNPs (5,663 parsimony-informative) from orthologous genomic segments (Table S2). This represents an average of only 1 SNP every ~500 bp across the entire genome and breadth of this species. A list of SNPs that define each branch and the homoplastic SNPs is provided in Table S3 to facilitate efforts by other researchers to place their strains in these established clades.

**Fig 1.**
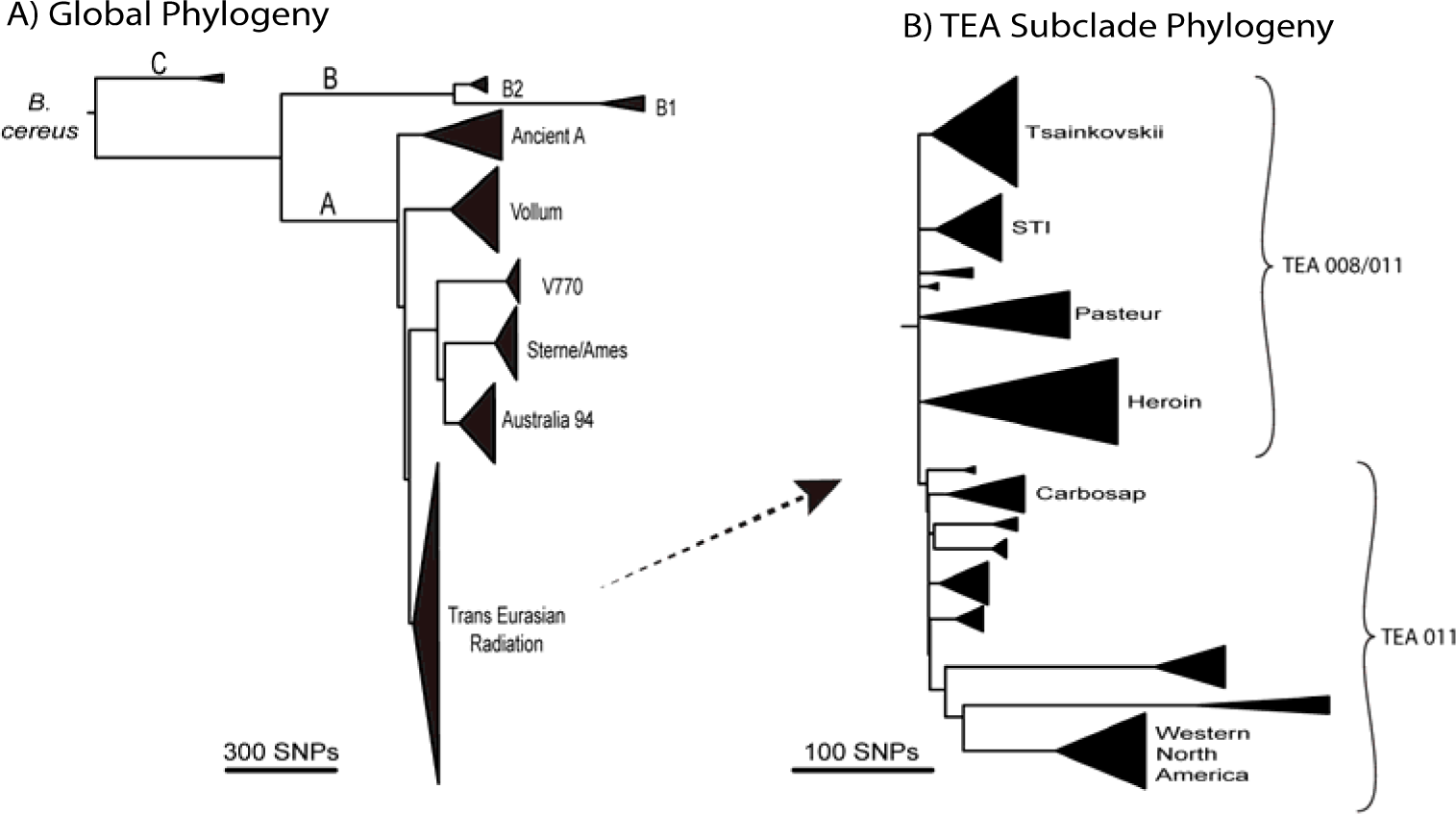
Phylogenetic Structure of *B. anthracis*. Core genome SNPs (11,989 total; 5,663 parsimony-informative) from whole genome sequences were analyzed by maximum parsimony to generate a phylogenetic tree. The major clades are collapsed in this figure but the complete tree is available in the supplemental material (Fig. S1). The overall consistency index is 0.98.

The deeper phylogenetic relationships (Fig. 1A) are consistent with those reported previously with a more limited number of genomes (Pearson, Busch et al. 2004, Van Ert, Easterday et al. 2007, Pearson, Okinaka et al. 2009, Marston, Allen et al. 2011, Khmaladze, Birdsell et al. 2014, Keim, Grunow et al. 2015, Pullan, Pearson et al. 2015, Vergnaud, Girault et al. 2016) as well as across different phylogenetic methods (Maximum Likelihood using the GTR model of evolution and Neighbor Joining). There are three major clades with C being basal to the A and B clades (Fig. 1A). Members of the A clade are most frequently observed across the globe (~90%) with B (~10%) and C (<1%) members being much less frequent (Van Ert et al. 2007). The A clade can be divided into four major monophyletic subclades with the “Ancient A” group being basal to the other subclades (Fig. 1A). Members of the TransEurAsia (TEA) subclade are most commonly observed as they have been highly successful across large and diverse geographic areas (Van Ert, Easterday et al. 2007).

The unusually short lengths of the deepest branches of the TEA clade, coupled with the high frequency of isolates and geographic expansion, is indicative of a rapid and extensive evolutionary radiation (Fig. 1B). Many sub-lineages of this clade diverged before mutations occurred, leading to a lack of synapomorphic characters (shared alleles that could group some of these sub-lineages together) and the existence of a large polytomy (a node with 7 immediate descendant lineages: Tsiankovskii, STI, Pasteur, Heroin, TEA 011, and two lineages with 1 and 2 genomes each). The expansion of each of these lineages also leads to multiple distinct groups, also often with very little topological resolution in the deeper nodes. Given the number of isolates assigned to the TEA 011 group, the TEA clade can be divided into two main subgroups: the paraphyletic TEA 008/011 (A.Br.008/011) and the monophyletic TEA (A.Br.011).

### Sverdlovsk Specimens Sequence Analysis

By direct DNA sequencing, we generated metagenomic data from paraffin-embedded formalin-fixed pathology specimens of two anthrax victims from the 1979 outbreak in Sverdlovsk USSR. The presence of *B. anthracis* DNA in these specimens had been previously established (Jackson, Hugh-Jones et al. 1998) and targeted gene sequencing had also been successful (Price, Hugh-Jones et al. 1999, Okinaka, Henrie et al. 2008); however, until recent technological advances in DNA sequencing, this could only be accomplished by first PCR amplifying small portions of the genome. Sequencing across both the MiSeq and HiSeq Illumina platforms produced ~300 million reads and 20 gigabases of nucleotide sequence data across both specimens. A direct mapping of reads against the finished genome of the Ames Ancestor genome with BWA-MEM demonstrated that only 1.2% of the total sequence data mapped to the reference genome. This is expected as DNA is from human tissue. The *B. anthracis* coverage represented an average sequencing depth of 24X across the chromosome, with >100X coverage of pXO1 and pXO2 plasmids. These data covered 99% of the Ames Ancestor genome, including both plasmids, with at least one read. Alignment stats are shown in Table 1.

**Table 1:** Alignment status for Sverdlovsk *B. anthracis* Genomes Table

| Library | Type | Length (bp) | Total Reads (pairs) | Trimmed reads (pairs) | Mapped reads (pairs) | Chr. coverage | pXO1 coverage | pXO2 coverage | Chr. breadth at 1x (%) |
| --- | --- | --- | --- | --- | --- | --- | --- | --- | --- |
| Sverd-1 | HiSeq | 93 | 6.4E+07 | 4.5E+07 | 5.4E+05 | 8x | 32x | 55x | 96% |
| Sverd-1 | MiSeq | 300 | 1.0E+07 | 3.4E+06 | 1.5E+05 | 1x | 3x | 27x | 30% |
| Sverd-2 | HiSeq | 93 | 7.5E+07 | 6.6E+07 | 1.0E+06 | 15x | 85x | 104x | 99% |
| Sverd-2 | MiSeq | 300 | 1.2E+07 | 3.6E+06 | 1.7E+05 | 1x | 6x | 28x | 36% |
| combined data: |  |  | 1.6E+08 | 1.2E+08 | 1.9E+06 | 25X | 126X | 214X | 99% |

From the reads, we assembled the Sverdlovsk genome into 128 contigs with an N50 size of 74Kb. A prediction of coding regions (CDSs) with Prodigal (Hyatt, Chen et al. 2010) on this assembly identified 5,579 CDSs; the same analysis on the Ames Ancestor genome identified 5,756 CDSs. This demonstrates that while most of the genome was successfully assembled, parts of the genome may have been dropped from the assembly, most likely from insufficient coverage or collapsed repeats.

### Data quality of the Sverdlovsk *B. anthracis* genome

Formalin fixation is known to damage nucleic acids and this was demonstrated by the small size of the extracted DNA fragments (Jackson, Hugh-Jones et al. 1998), but its effect upon the validity of the Sverdlovsk genomic sequence was unknown. The intrinsic error rate in a sequencing project can be measured by mapping individual sequencing reads to a high quality reference genome. This generates an estimate of the raw read error rate at each nucleotide and across the whole genome, representing a sequencing quality measurement particularly relevant to SNP identification. In a comparison of *B. anthracis* sequencing reads from Sverdlovsk pathology specimens to those from DNA isolated from culture, we observe a higher number of errors (Fig. S1). The average rate per nucleotide was 0.2% for the culture generated DNA versus 0.5% for the formalin fixed tissue. In both cases, a true polymorphism would not be determined from a single read but rather from the consensus of multiple read coverage at any particular genomic position; however, see Sahl et al (Sahl, Schupp et al. 2015) for a low coverage SNP calling strategy. We further examined the consequences of this differential error rate by searching for the conservation of known SNPs along a particular phylogenetic path within these genomes. These were identified in the 193 genome phylogeny (Fig. 1), independent of the Sverdlovsk genome. There were 329 known SNP changes along the branches that connect the Ames Ancestor reference to the composite Sverdlovsk genome (Figure 1 and Table S3; Supplemental figures S2, S6 and S9). All 329 SNP sites were present in the composite genome assembly. Excluding 29 SNP sites on the pXO1 and pXO2 plasmids because they have higher copy numbers, the coverage per SNP averaged 20X at 273 of the remaining 300 genomic positions on the chromosome. Fourteen of the other chromosomal SNP sites contained less than 10 reads per site but still corresponded exactly to the expected base changes. Overall, we were able to discover and verify all of the known SNPs using the Sverdlovsk pathology specimen sequencing data. Based upon these two error estimations, we are confident that the sequenced genomes are of sufficient quality to justify our conclusions.

### Phylogenetic Position of the Sverdlovsk Strain

Based upon shared SNPs, the Sverdlovsk genomes fall within the “Tsiankovskii” subclade of the TEA 008/011 group (Figure 1B). Within this group, it is most closely related to two other Asian strains both of which are used as vaccines. There are only 13 SNPs on the branch to the Sverdlovsk genomes, 25 on the branch to Tsiankovskii, and 52 on the branch to Cvac02 (Tables S2-S4). These three genomes emerge from a polytomy, showing rapid divergence of these lineages before shared SNPs could arise. As this clade is comprised of laboratory strains, this divergence may be due to anthropogenic establishment of different lineages from a laboratory stock. Other clade members were isolated from anthrax-killed animals and are mostly Eastern European in origin, with the exception of one from China and one from Norway. Therefore, with the exception of the three “domesticated” strains, the clade members are naturally occurring wild type strains.

### The Sverdlovsk *B. anthracis* genome specific SNPs

The sequencing and analysis of Sverdlovsk genomes offers an opportunity to detect SNPs and to look for possible strain mixtures or contaminating DNA profiles from two of the tissue samples. To do this, nucleotides from individual reads are tabulated and less than 100% agreement represents potential errors or mixtures at that genomic position. In particular, we are interested in the 13 SNPs that are unique to Sverdlovsk genomes as they allow a comparison to all other strains outside this group to identify mixtures. Table 2 shows the consensus read results from Sverdlovsk specific SNPs and overall there are only 7 variants, resulting in an error rate of 1.6%, which is only slightly higher than the overall error rate of 0.5%. In addition, we note that 6 of the 7 differences are located near the ends of reads where the error rate is higher (data not presented). One SNP (NC_007530:5138018) was detected between the two specimens and this contrast appears to represent a real difference as it was supported by >18 reads. A small number of SNPs between these two specimens might be observed given the population size associated with large-scale production and subsequent amplification *in vivo*. Otherwise, we find no evidence in these two particular Sverdlovsk specimens for strain mixtures. It is important to recognize that these two specimens did not show mixed alleles at the *vrrA* locus analyzed by Jackson et al. (Jackson, Hugh-Jones et al. 1998).

**Table 2.** Read Mixtures at Sverdlovsk Genome Specific SNPs

| SNP Site | SNP | Read Depth | % Consensus | Genomic Element |
| --- | --- | --- | --- | --- |
| 28576 | G to A | 78 | 97.2 | pX02 |
| 59151 | C to T | 49 | 100 | Chromosome |
| 112138 | A to G | 36 | 100 | Chromosome |
| 602999 | C to A | 34 | 100 | Chromosome |
| 1359179 | C to T | 27 | 100 | Chromosome |
| 2138718 | A to T | 20 | 100 | Chromosome |
| 2979549 | A to G | 27 | 100 | Chromosome |
| 3593664 | T to C | 31 | 89.9 | Chromosome |
| 4034596 | G to A | 17 | 93.7 | Chromosome |
| 4236707 | T to C | 30 | 100 | Chromosome |
| 4504222 | C to T | 22 | 100 | Chromosome |
| 4896833 | T to C | 17 | 100 | Chromosome |
| 5186004 | C to A | 46 | 100 | Chromosome |

### Genetic Engineering Evidence

Particular genes and SNP signatures in the Sverdlovsk genomes were examined for evidence of genetic manipulation of this strain. In the chromosome, fluoroquinolone resistance is known to be determined by amino acid changes in the *gyrA* and *parC* genes (Price, Vogler et al. 2003), rifampicin resistance is associated with changes in the *rpoB* gene (Vogler, Busch et al. 2002), and penicillin resistance is associated with changes in β-lactamase gene expression (Ross, Thomason et al. 2009). With regards to amino acid changes in associated genes, the Sverdlovsk genomes contained wild type drug susceptible alleles. The cereolysin genes and plasmid sequences used by Russian scientists to alter *B. anthracis* phenotypes (Stepanov, Marinin et al. 1996, Pomerantsev, Staritsin et al. 1997) were not present. In addition, the read data were examined for other common genetic engineering vectors, which were not detected, from an alignment of raw reads against the NCBI UniVec database. The alignment of the 128 contigs to the Ames Ancestor revealed no novel genes (Fig. 2), though this was not a closed genome. Hence, there is no evidence from this analysis of either molecular-based genetic engineering or classical bacteriological selection for altered drug resistance phenotypes.

**Figure 2.**
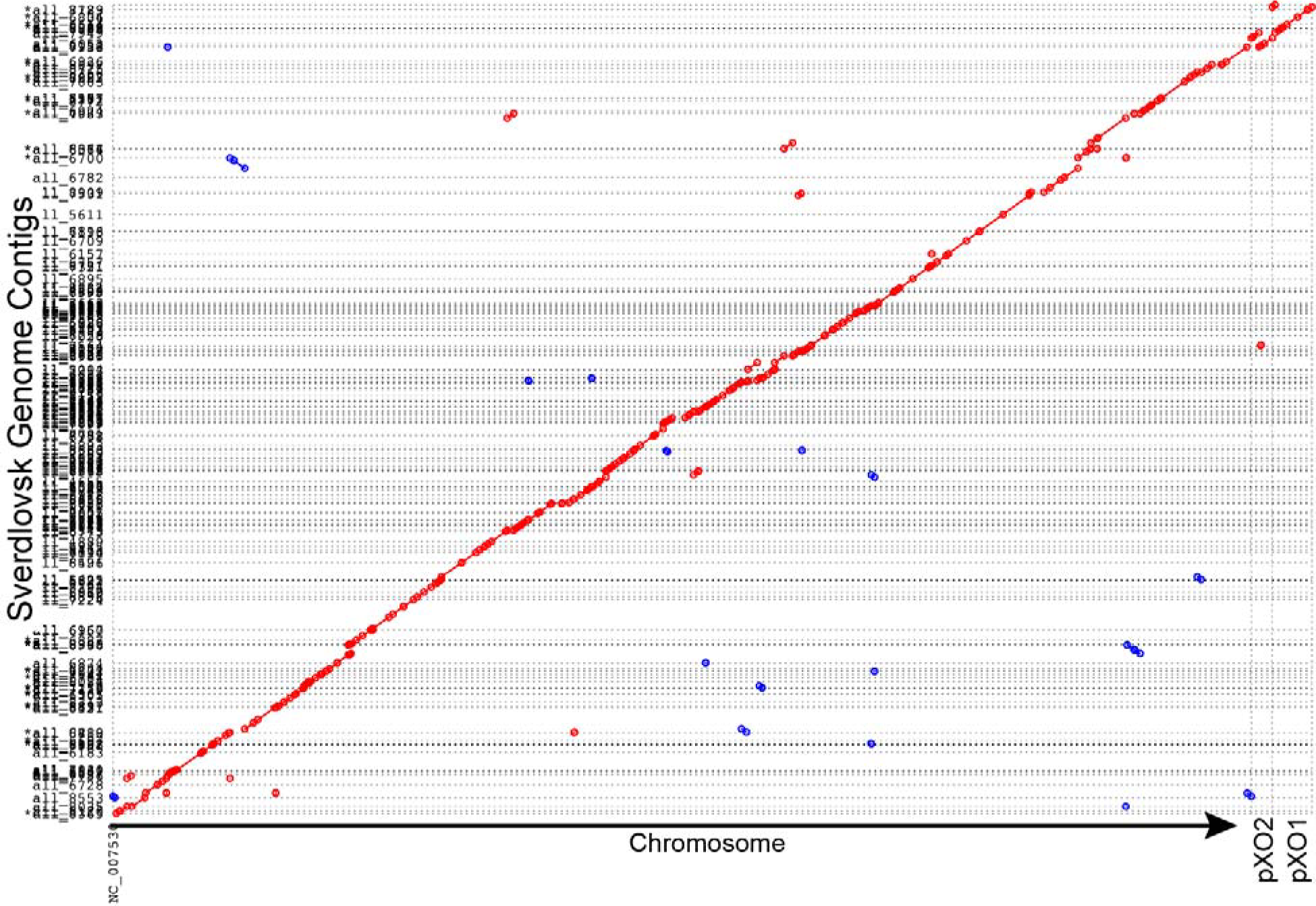
Sverdlovsk Contigs aligned to the Ames Ancestor Genome. The reads from both autopsy specimens were combined for de novo assembly, which resulted in 128 contigs. These are aligned against the Ames Ancestor chromosome and two plasmids and the synteny was visualized with mummerplot. Greater than 99% the Ames Ancestor genome is represented by these 128 contigs.

**Figure 3.**
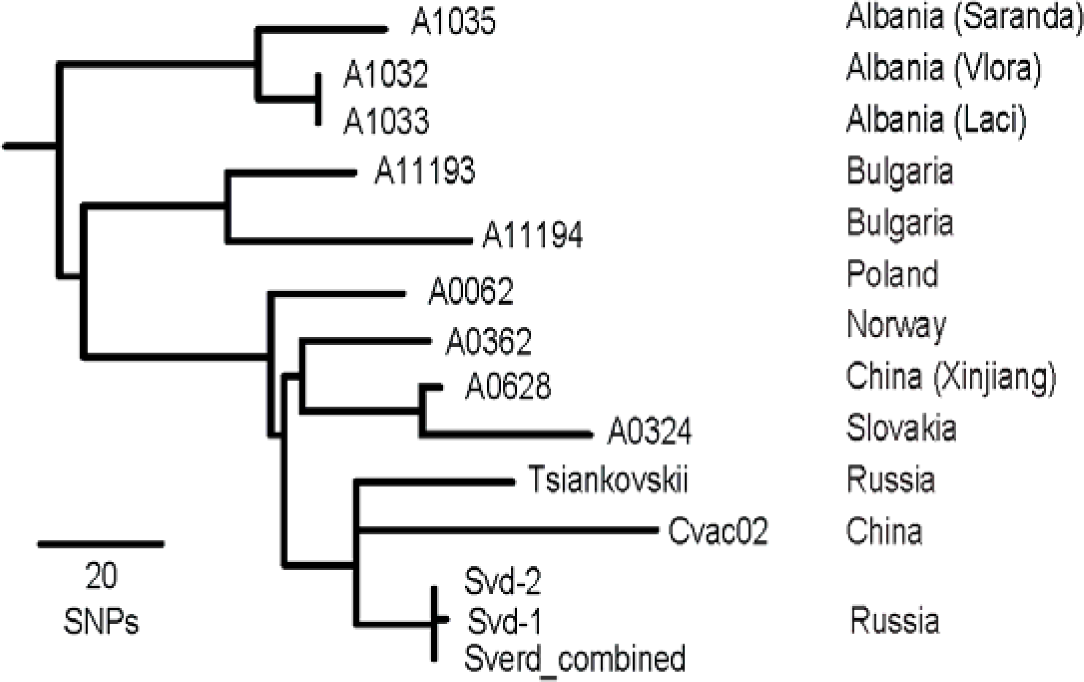
The Tsiankovskii clade. A phylogenetic tree of the closest relatives to the Sverdlovsk genomes are shown. One SNP was discovered between Svd-1 and Svd-2. The Sverd combined genotype is identical to Svd-2.

## Discussion

The *B. anthracis* global phylogeny is one of the most robust evolutionary reconstructions available for any species. This is possible because core genome SNPs represent highly stable evolutionary characters with very low homoplasy and their rarity in this genome precludes any effects from mutational saturation. This species' evolutionary reconstruction is a function of its spore-vegetative cycle biology and in particular, its ecological niche. The dormant spore stage is important for its dispersal, transmission, limiting evolutionary changes and restricting interactions with near neighbor *Bacillus* species, making it resistant to horizontal gene transfer. Hence, the *B. anthracis* pan-genome is only slightly larger than the core genome, with variation primarily due to decay via gene deletion. Environmental growth outside the host is possible, but does not appear to represent a significant opportunity to shape this bacterium’s genome and evolution. Long quiescent periods in the spore phase may create a “time capsule” where few or no mutations are generated, which has resulted in a highly homogeneous pathogen. In this sense, its niche differs from its close relative *B. cereus*, which is environmentally adapted with occasional pathogenic replication in a host (Zwick, Joseph et al. 2012). Fortuitously, the genome variation that we can identify through whole genome sequencing generates insights into anthrax history and allows predictions about its ecology.

The clade structure we observe with whole genome sequencing is consistent with previous descriptions using lower resolution methods or few genome sequences. What we add in this report is the precise definition of branching points, accurate branch length determinations, and the definition of canonical evolutionary characters for strain identification. Branch topology determination has been problematic with other molecular methods because of the abundance of short branches and polytomies at critical positions in the evolutionary structure. The A-clade
itself, but in particular its subclade TEA, are evidence for evolutionary radiations representing genetic bottlenecks, long-distance dispersal and bursts in the fitness of these lineages. Even in a radiation, binary fission of replicating bacterial cells should result in phylogenetic structure that could be identified with sufficiently discriminatory methods. But in some cases, such as with the TEA clade, even whole genome analysis does not yield topological phylogenetic structure, arguing for a very tight genetic expansion. This subclade contains a large portion of the world’s anthrax burden (Van Ert, Easterday et al. 2007), making this radiation event seminal. Molecular clock analyses for 106 sub-root dated isolates (Table S1 and Fig. S13) and the 48 dated TEA isolates (Fig. S14) have revealed a complete lack of temporal signal among this relatively contemporary dataset, leaving the exact timing of this radiation dependent upon phylogeographic hypotheses. These models are controversial and vary widely in their temporal predictions (Kenefic, Pearson et al. 2009, Vergnaud, Girault et al. 2016). To insure that the lack of molecular clock signal is not due to error arising from various sequencing methods, we pruned the phylogeny to clade A isolates with sister taxa that have dates of isolation within 5 years of each other. We then removed all non-parsimony informative sites, such that only shared SNPs (aside from a small number of homoplastic SNPs) were used to reconstruct the phylogeny as we assume that sequencing errors are unlikely to occur on shared branches. As in the former root-to-tip analyses, a temporal signal was not evident (Fig. S15). Ancient genomes from archeological sites would greatly assist in the temporal calibration of key branch points.

Detailed genome databases are a great resource for public health and forensic investigations of disease outbreaks (Aarestrup, Brown et al. 2012). As disease events occur, they allow for the real time matching of similar types and source identification. But pathogens are dynamic and databases must be continually updated with isolates from contemporary outbreaks. For some pathogens, a few months can allow for genomic divergence that will make source tracking problematic (Hendriksen, Price et al. 2011, Eppinger, Pearson et al. 2014). The availability of high quality reference databases set the stage for further sampling (Keim, Grunow et al. 2015). It is important to define the relevant subpopulation for additional investigative sampling (Keim 2011) and this will not be possible prior to a disease outbreak.

Inspired by other preserved pathology tissue DNA analyses (Devault, Golding et al. 2014), two *B. anthracis* genome sequences from victims of the Soviet military accident in Sverdlovsk Russia were generated by deeply sequencing formalin fixed autopsy specimens. Although only ~1.2% of the sequenced reads were associated with the pathogen, enough information was obtained for high-resolution phylogenetics and for draft genome assemblies. A higher than normal error rate was observed in the Sverdlovsk samples, likely due to the nature of the specimen preservation, but sufficient depth of coverage was still obtained to accurately genotype known SNP loci and to identify strain specific polymorphisms. Contigs assembled from the reads are syntenic with reference genomes and consistent with isolates from natural anthrax outbreaks with no extraneous reads associated with cloning vectors or novel toxins. Additionally, there was no evidence of *B. anthracis* strain mixtures in these two particular specimens. Jackson et al. (Jackson, Hugh-Jones et al. 1998) reported mixed alleles at the *vrrA* locus for some tissue samples, but not the two analyzed in this report. The vrrA locus could not be assembled from these specimens due its repeat structure and the other victim specimens had very limited DNA that was prohibitive of metagenomic analysis. Hence, our analysis does not eliminate the possibility that mixed strains were involved in the Sverdlovsk anthrax outbreak.

The Soviet “battle strain” 836 was isolated from nature (Alibek and Handelman 1999) and used for industrial spore production in the 1960’s and 70’s, which was mostly prior to the advent of recombinant DNA methods. Traditional selection for mutants resistant to antibiotic resistance was certainly possible prior 1979, but no such mutations are evident in the Sverdlovsk strain genomes. The great similarity of the genomes to other natural isolates argues for minimal laboratory manipulation. It is well established that *B. anthracis* attenuates with laboratory culturing and selection for drug resistance frequently has secondary phenotypic consequences that would not be desirable for a weapons strain (Price, Vogler et al. 2003). All of this is highly suggestive of a weapons program that identified a suitable strain, maintained master cell stocks to avoid extensive passage and performed minimal manipulations in order to maintain virulence. This strategy must have been used to produce large quantities of highly virulent material as evidence by the anthrax deaths in 1979.

## Acknowledgements

We authors would like to thank three reviewers who provided critical and constructive comments of the penultimate manuscript: Matt Meselson, Tim Read and Nick Loman. This work was supported with a contract (HSHQDC-15-C-B0068) from the Department of Homeland Security Science and Technology Directorate.

---

Table S1: List of *B. anthracis* strains, genome accession numbers, and associated metadata.

Table S2: SNP character states for 11,989 SNPs across all 193 *B. anthracis* genomes.

Table S3: Branch assignments for all SNPs.

Table S4: SNP character states for the 376 SNPs within the Tsiankovskii subclade.

**Figure S1.**
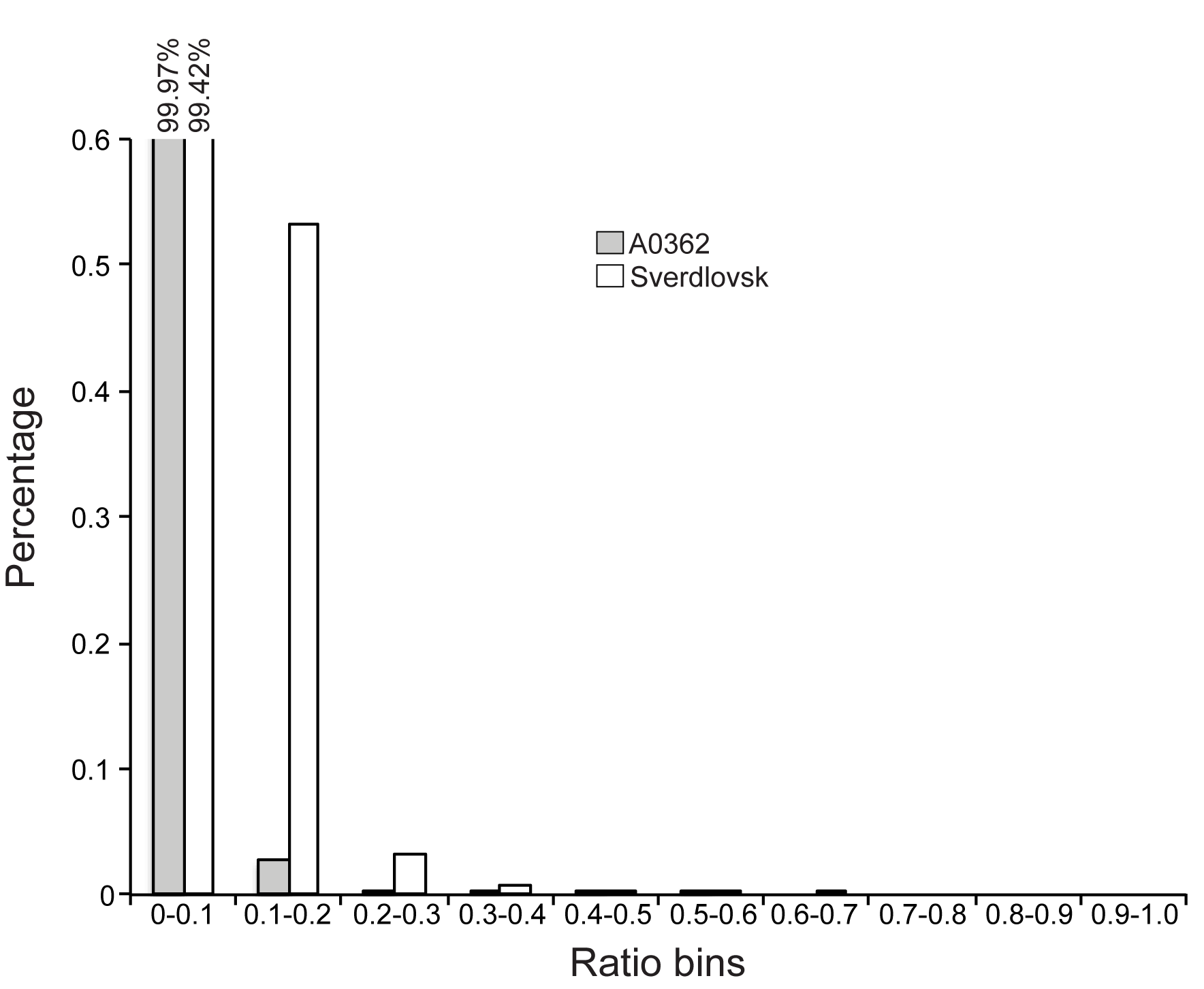
Read error rate profile across the genome for Sverd and a culture DNA (A0362: SRR2968203). Reads were aligned to Ames Ancestor and the composition of base calls was compared. Error rates were determined by dividing the number of minor allele calls by the total number of calls. The error rates were then binned into categories from no error to total error. The frequency of calls in each bin are represented by the height of histograms. The results demonstrate that while both genomes had low error, the Sverdlovsk genome had a higher error profile than a contemporary, pure culture.

**Figure S2:**
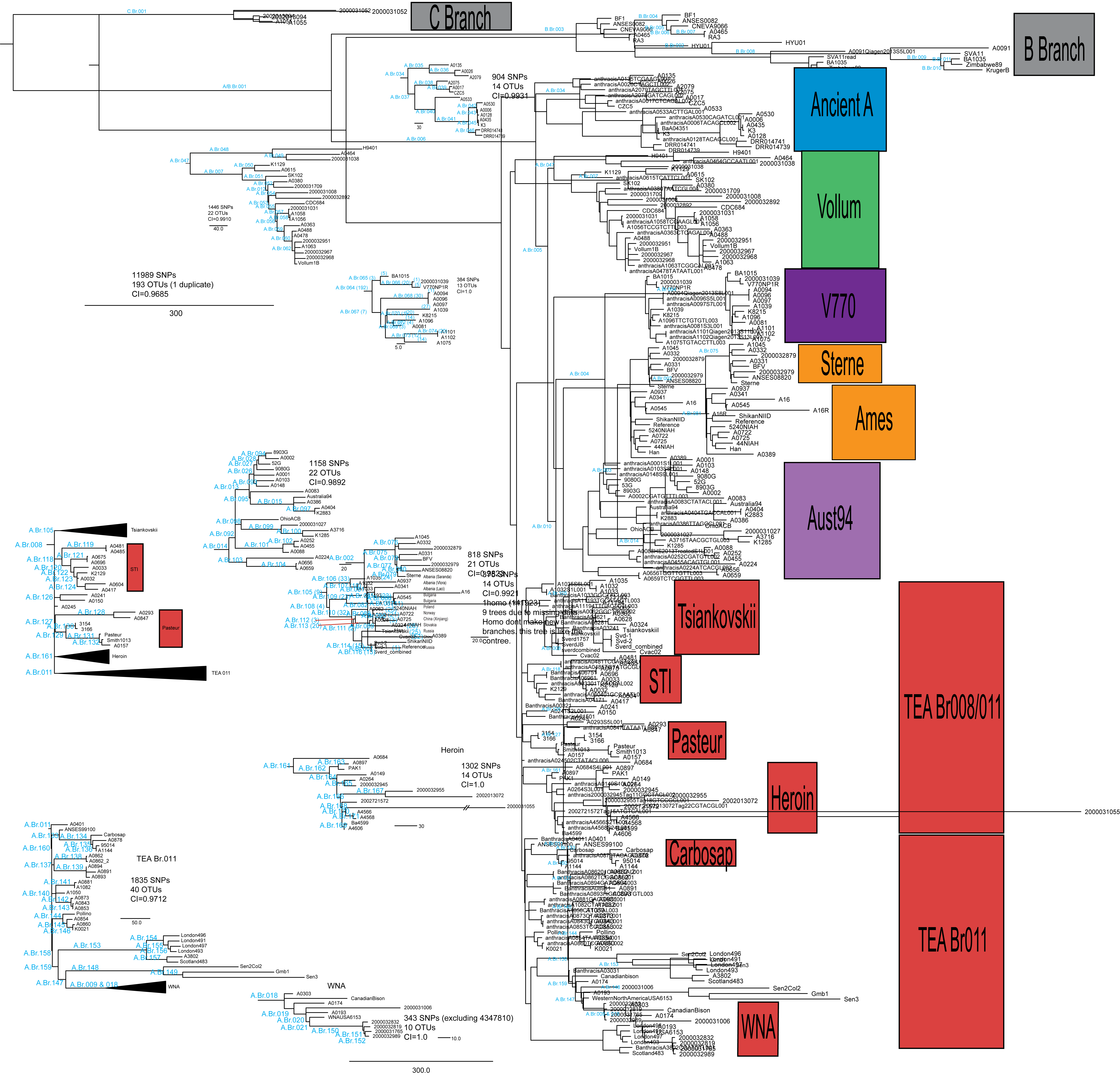
Maximum parsimony phylogeny of 193 *B. anthracis* genomes. CI (excluding parsimony uninformative characters) = 0.9657. Names of major branches are indicated in blue text. Branch names within each clade are included in supplemental figures dedicated to each clade.

**Figure S3:**
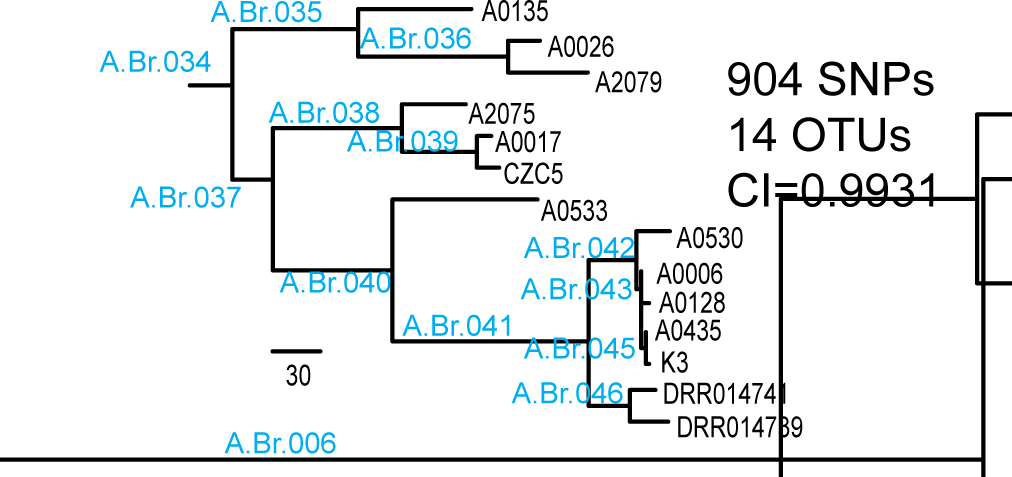
Maximum parsimony phylogeny of the “Ancient A” clade. The formal name for this clade is A.Br.006/005. This clade currently contains 14 genomes and 904 SNPs. CI (excluding parsimony uninformative characters) = 0.9931. Names of branches are indicated in blue text.

**Figure S4:**
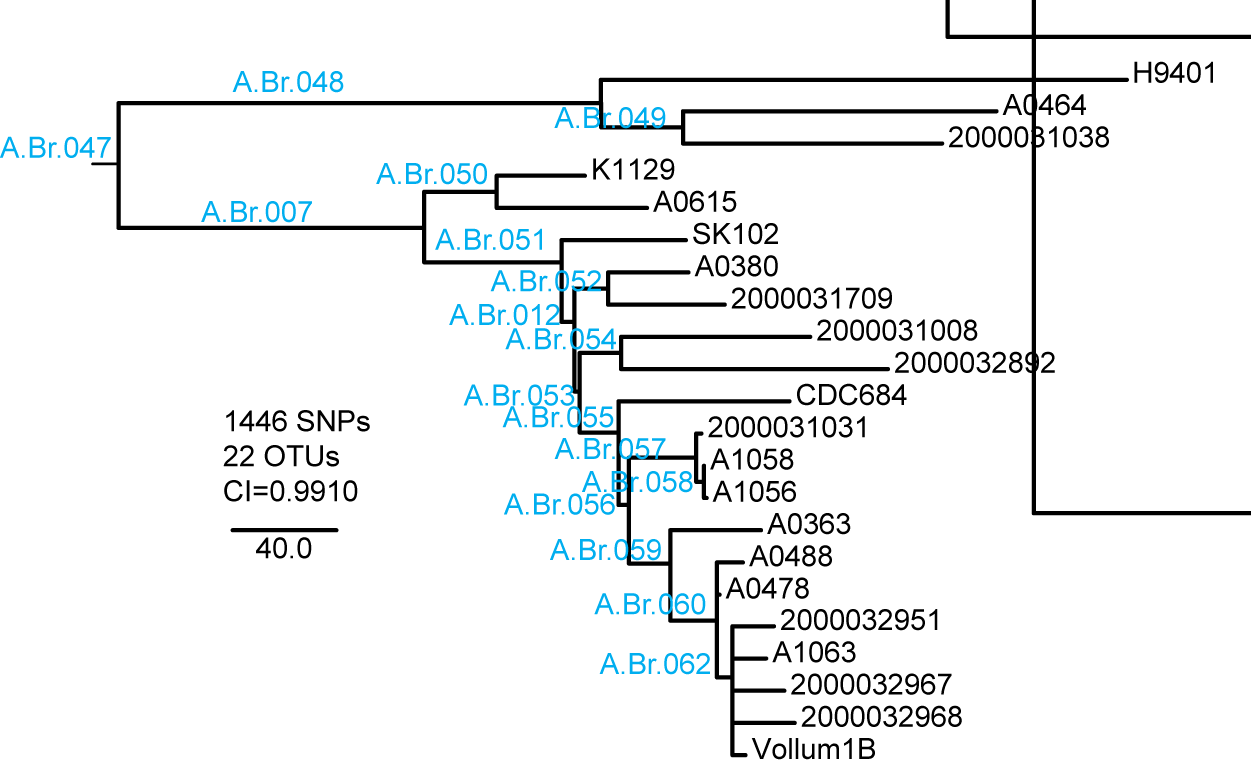
Maximum parsimony phylogeny of the “Vollum” clade. The formal name for this clade is A.Br.005/010. This clade currently contains 22 genomes and 1,446 SNPs. CI (excluding parsimony uninformative characters) = 0.9910. Names of branches are indicated in blue text.

**Figure S5:**
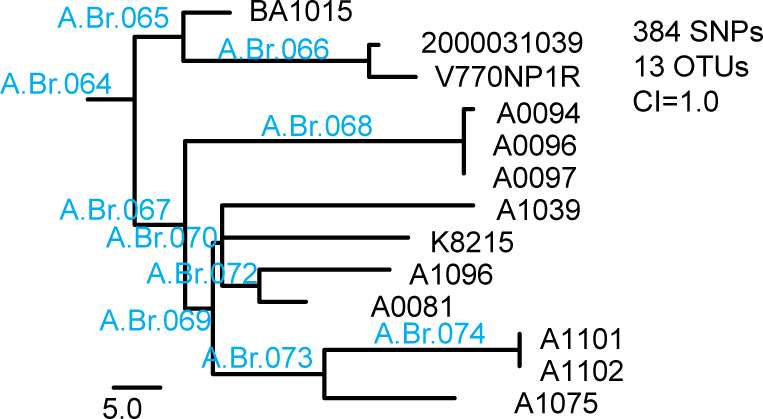
Maximum parsimony phylogeny of the “V770” clade. The formal name for this clade is A.Br.004/003. This clade currently contains 13 genomes and 384 SNPs. CI (excluding parsimony uninformative characters) = 1.0. Names of branches are indicated in blue text.

**Figure S6:**
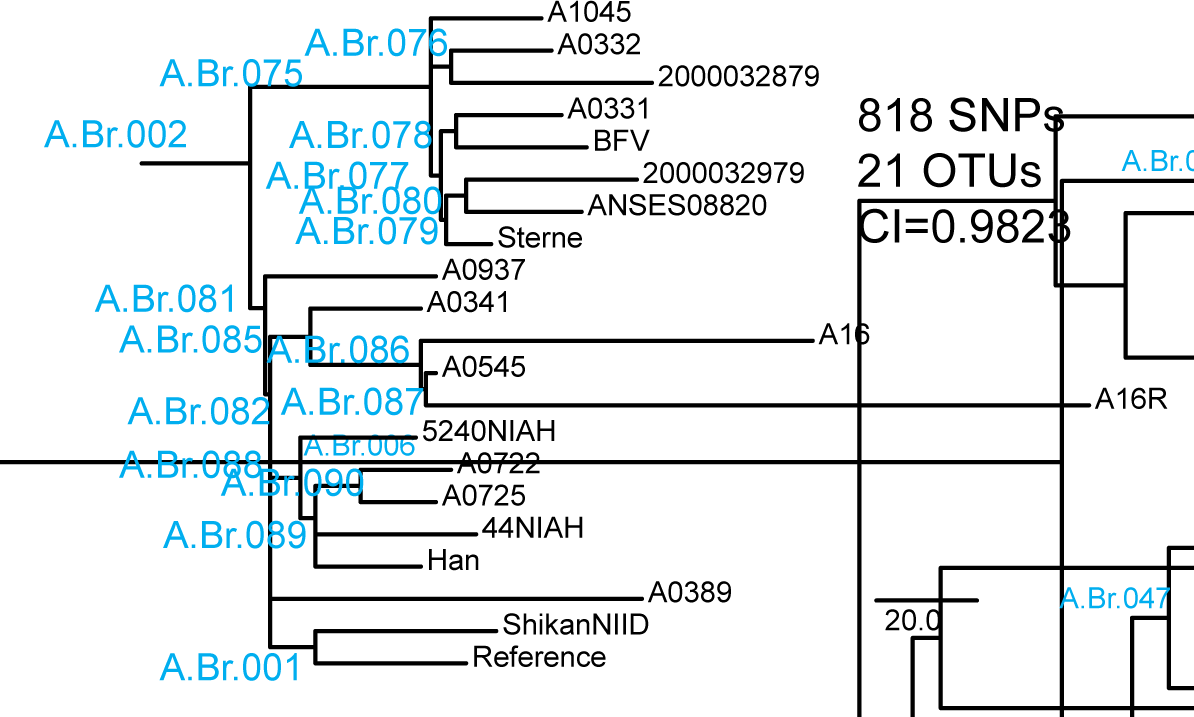
Maximum parsimony phylogeny of the “Sterne/Ames” clade. The formal name for this clade is A.Br.003/014. This clade currently contains 21 genomes and 818 SNPs. CI (excluding parsimony uninformative characters) = 0.9823. Names of branches are indicated in blue text.

**Figure S7:**
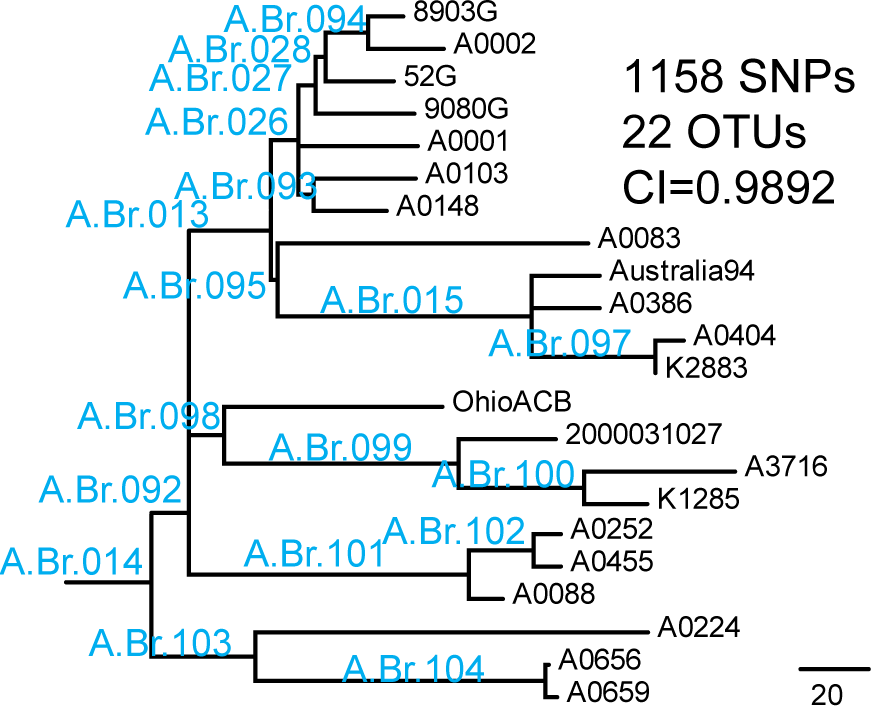
Maximum parsimony phylogeny of the “Aust94” clade. The formal name for this clade is A.Br.003/002. This clade currently contains 22 genomes and 1,158 SNPs. CI (excluding parsimony uninformative characters) = 0.9892. Names of branches are indicated in blue text.

**Figure S8:**
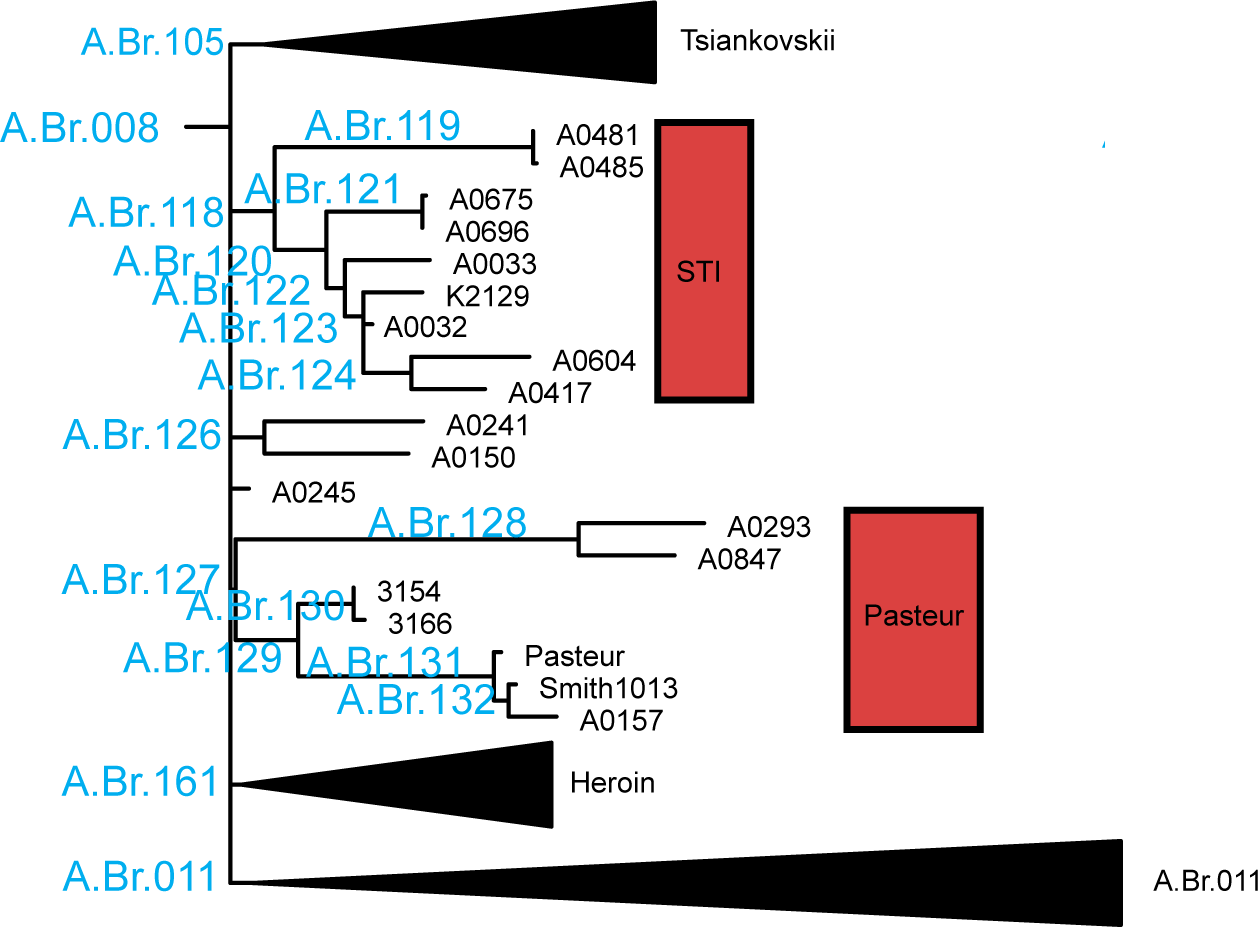
Phylogeny of the “TEA” clade. This clade contains many large subclades that are presented in detail in supplemental figures S9-S11. Names of major branches are indicated in blue text.

**Figure S9:**
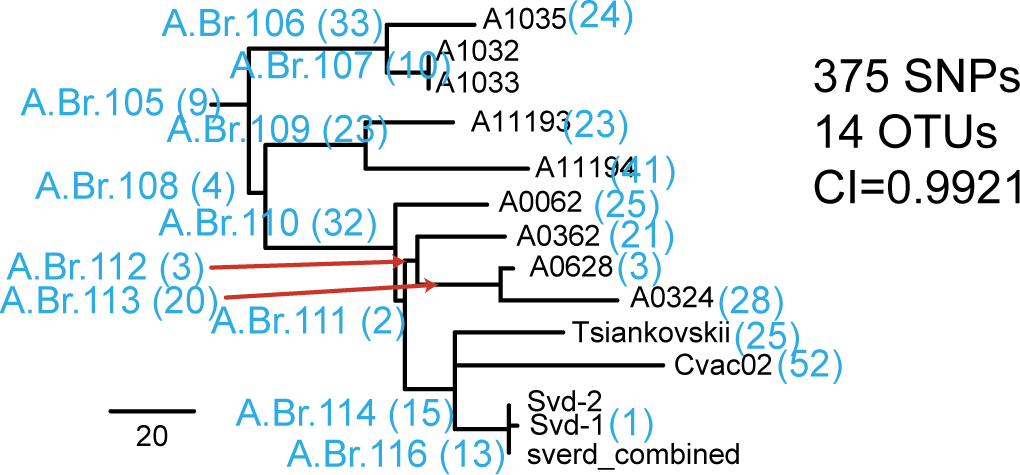
Maximum parsimony phylogeny of the “Tsiankovskii” subclade (see also Fig. 3). This subclade is part of the “TEA” clade and is within the A.Br.008/011 clade. This subclade currently contains 14 genomes and 375 SNPs. CI (excluding parsimony uninformative characters) = 0.9921. Names of branches and branch lengths are indicated in blue text.

**Figure S10:**
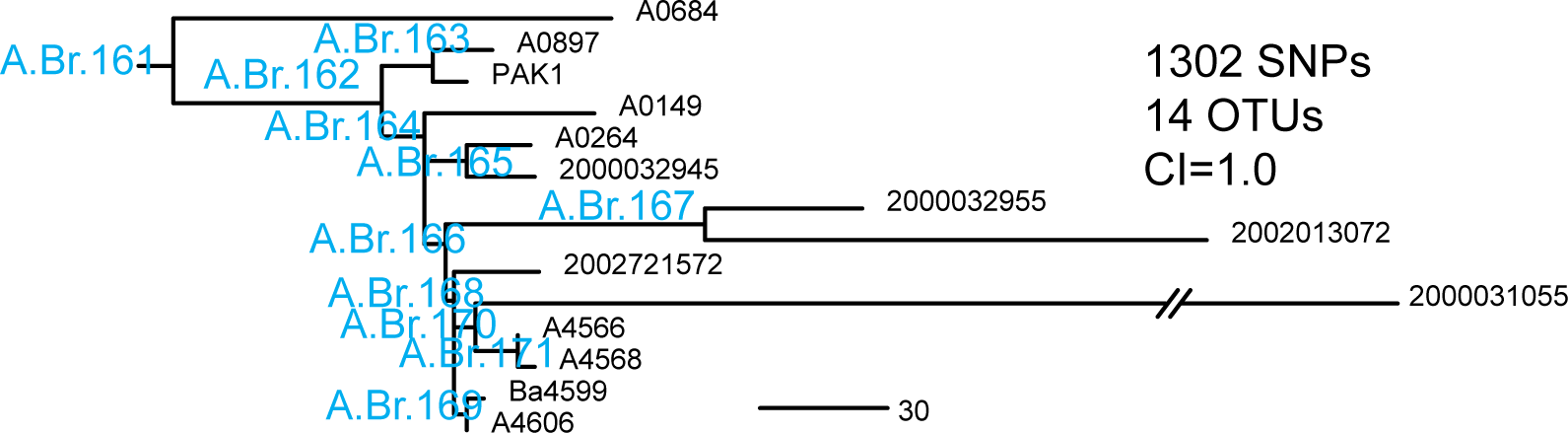
Maximum parsimony phylogeny of the “Heroin” subclade. This subclade is part of the “TEA” clade and is within the A.Br.008/011 clade. This subclade currently contains 14 genomes and 1,392 SNPs. CI (excluding parsimony uninformative characters) = 1.0. Names of branches are indicated in blue text.

**Figure S11:**
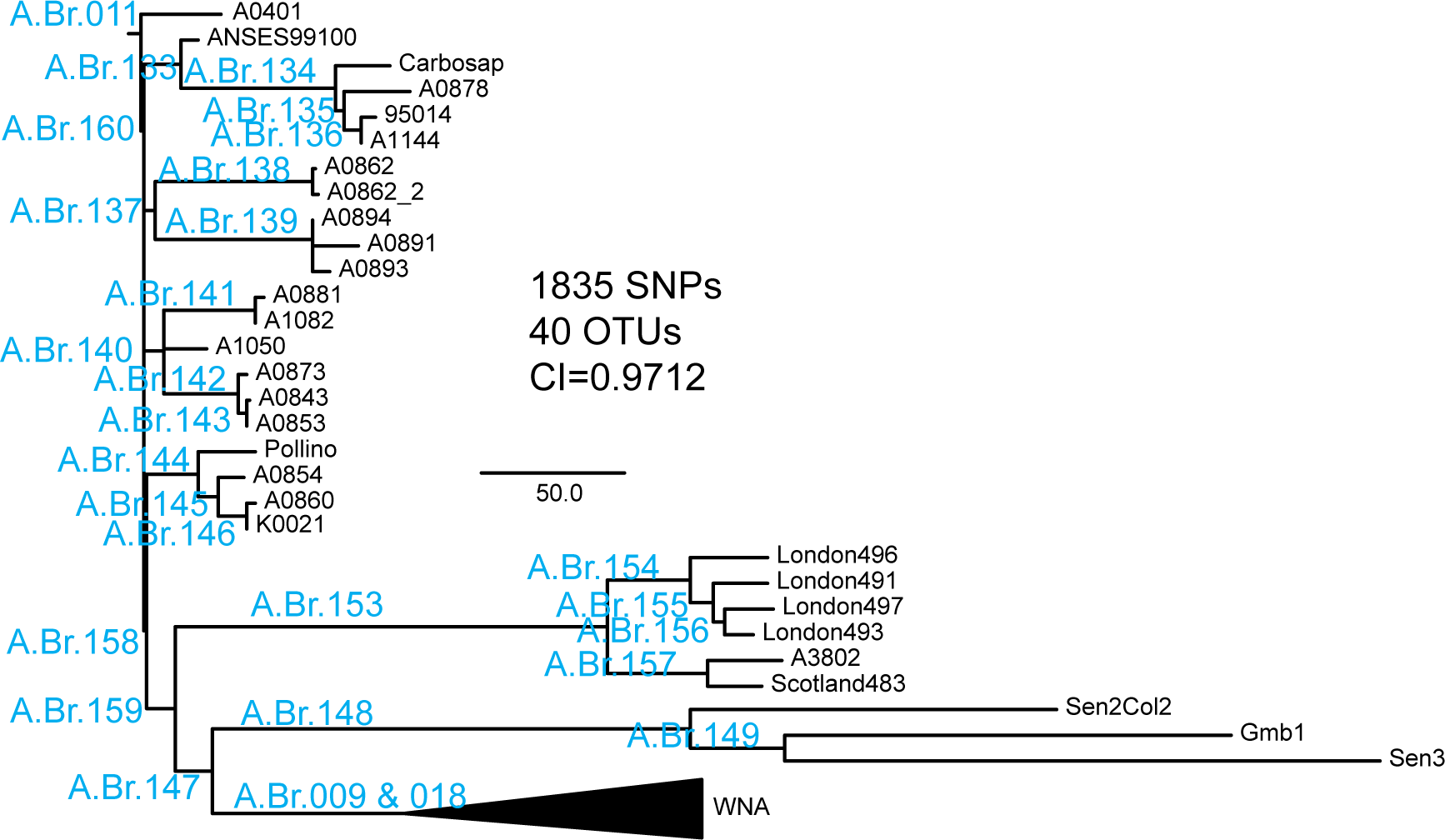
Phylogeny of the “TEA 011” subclade. This subclade is part of the “TEA” clade. This clade contains the “WNA” subclade that is presented in detail in supplemental figure S11. This subclade currently contains 40 genomes and 1,835 SNPs. CI (excluding parsimony uninformative characters) = 0.9712. Names of branches are indicated in blue text.

**Figure S12:**
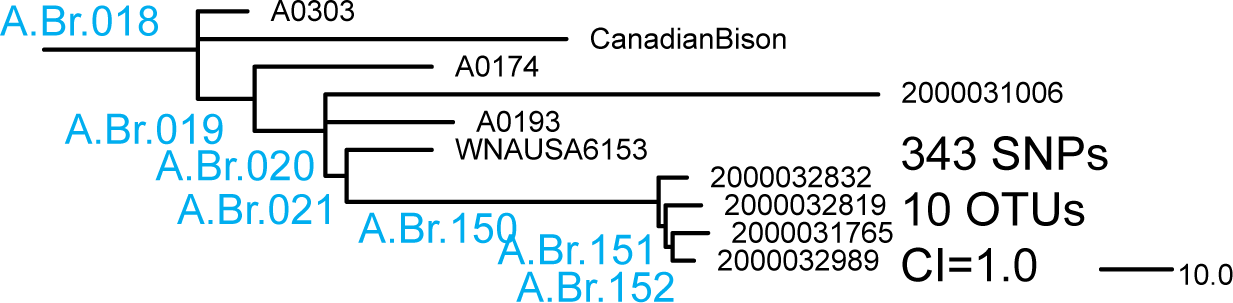
Phylogeny of the “WNA” subclade. This subclade is part of the “TEA” clade. This subclade currently contains 10 genomes and 343 SNPs. CI (excluding parsimony uninformative characters) = 1.0. Names of major branches are indicated in blue text.

**Figure S13:**
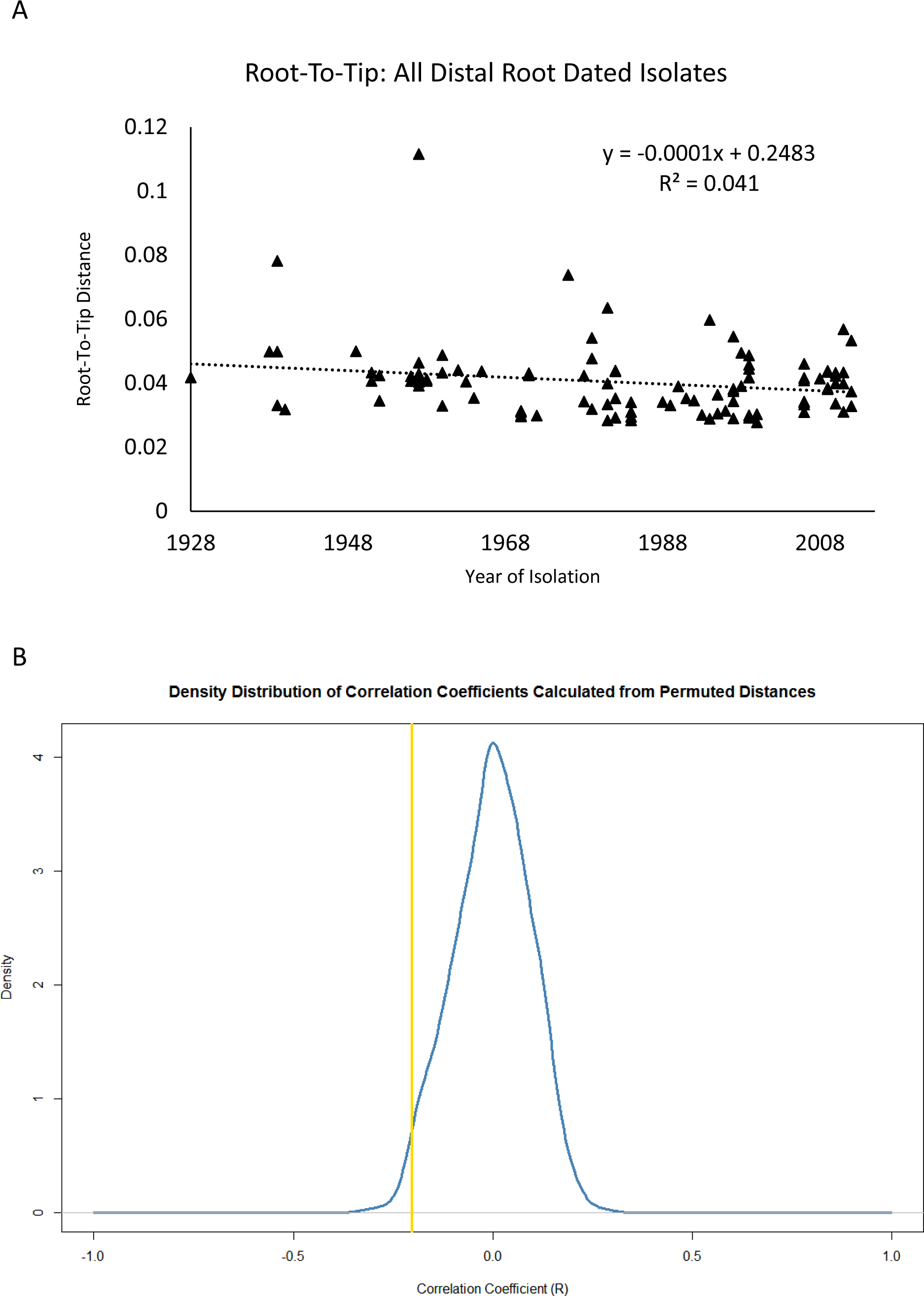
Molecular clock analysis for all genomes with isolation dates, except for the three root-most isolates (2002013094, A1055, and 2000031052). A) Linear regression analysis of root-to-tip distances extracted by TempEst (Rambaut, Lam et al. 2016) from a neighbor joining tree reconstructed in MEGA7 (Kumar, Stecher et al. 2016). The negative slope and low R_2_ value indicates that time does not explain root-to-tip distances, measured in substitutions per site. B) A permutation test was conducted, where dates were randomly shuffled among the root-to-tip distances 1000 times, and each time a linear regression was conducted. The observed Correlation Coefficient (r=−0.2, yellow line), was plotted among the distribution of r values from the permutations. The observed r value (yellow line) is greater than only 19 of 1000 values composing the distribution. Additionally, the negative r value indicates that the relationship is root-to-tip distance is not correlated with time.

**Figure S14:**
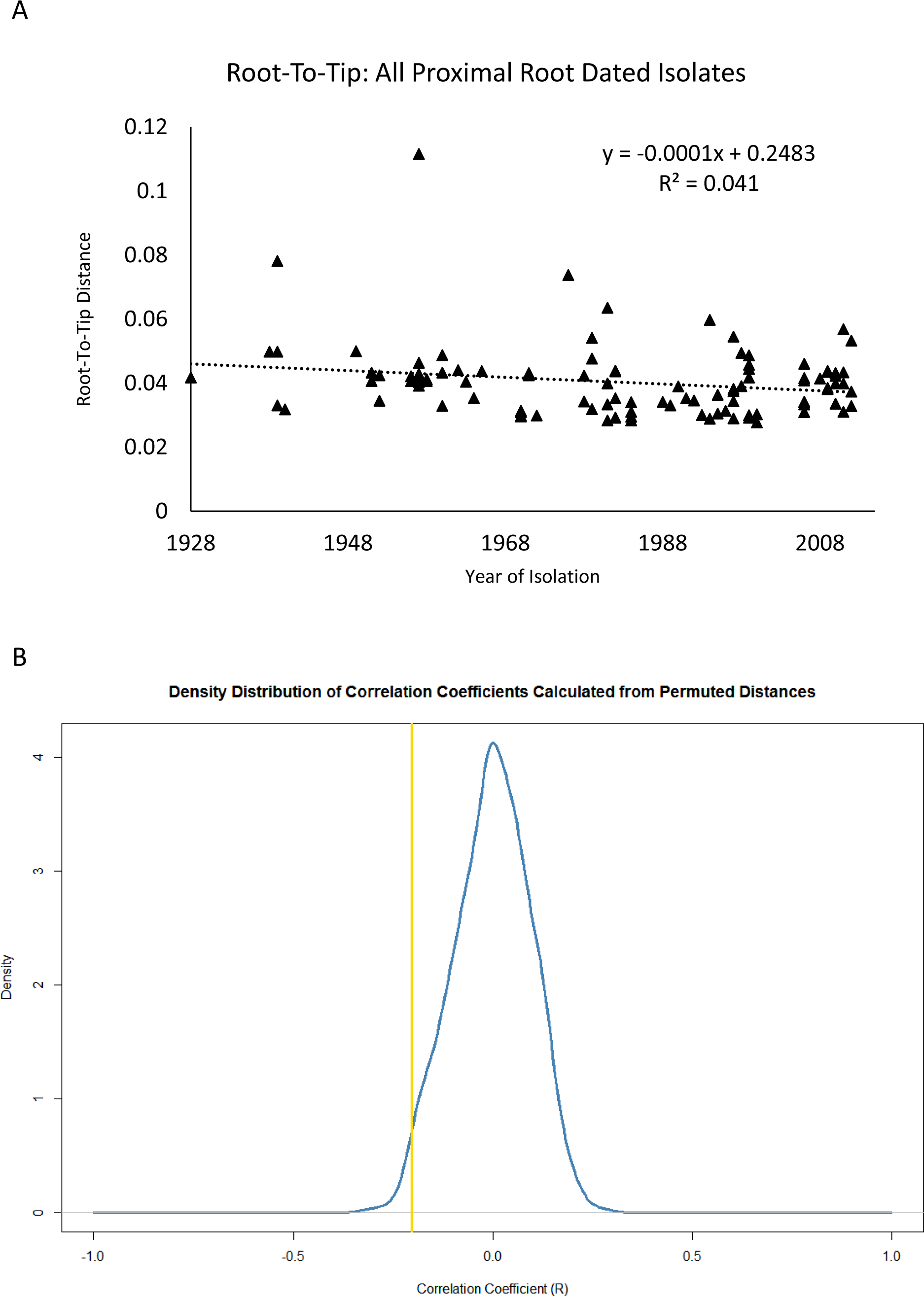
Molecular clock analysis for genomes in the TEA clade, except for the hypermutator isolate (2000031055). A) Linear regression analysis of root-to-tip distances extracted by TempEst (Kumar, Stecher et al. 2016) from a neighbor joining tree reconstructed in MEGA7 (Kumar, Stecher et al. 2016). The nearly horizontal slope and weak correlation (low R^2^ value) indicates that time does not explain root-to-tip distances, measured as substitutions per site. B) A permutation test was conducted, where dates were randomly shuffled among the root-to-tip distances 1000 times, and each time a linear regression was conducted. The observed Correlation Coefficient (r= 0.03, yellow line) value, was plotted among the distribution of r values from the permutations. The observed r value (yellow line) is greater than 651 of 1000 values composing the distribution, indicating that the Correlation Coefficient is no greater than expected by chance.

**Figure S15:**
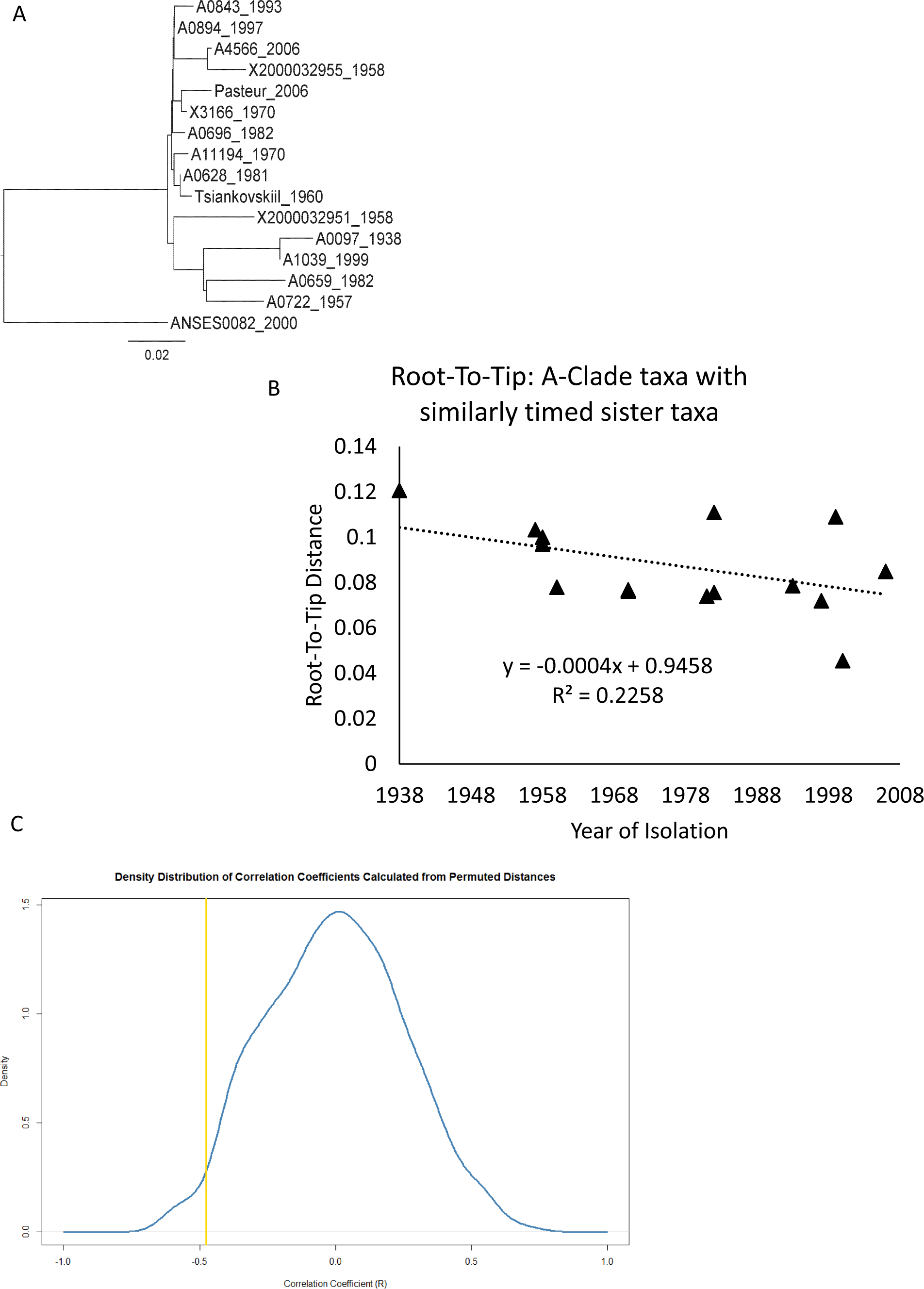
Molecular clock analysis using only parsimony informative SNPs for A-clade (ingroup) genomes with at least one sister taxon dated within five years. A) Neighbor-joining tree including remaining taxa. B) Linear regression analysis of root-to-tip distances extracted by TempEst (Kumar, Stecher et al. 2016) from a neighbor joining tree reconstructed in MEGA7 (Kumar, Stecher et al. 2016). The negatively correlated slope indicates that time does not explain root-to-tip distances, measured as substitutions per site. C) A permutation test was conducted, where dates were randomly shuffled among the root-to-tip distances 1000 times, and each time a linear regression was conducted. The observed Correlation Coefficient (r= -0.47, yellow line) value, was plotted among the distribution of r values from the permutations. The observed r value (yellow line) is greater than 22 of 1000 values composing the distribution, indicating that the Correlation Coefficient is no greater than expected by chance.

## References

Aarestrup, F. M., E. W. Brown, C. Detter, P. Gerner-Smidt, M. W. Gilmour, D. Harmsen, R. S. Hendriksen, R. Hewson, D. L. Heymann, K. Johansson, K. Ijaz, P. S. Keim, M. Koopmans, A. Kroneman, D. Lo Fo Wong, O. Lund, D. Palm, P. Sawanpanyalert, J. Sobel and J. Schlundt (2012). “Integrating genome-based informatics to modernize global disease monitoring, information sharing, and response.” Emerg Infect Dis 18(11): e1.

Abramova, F. A., L. M. Grinberg, O. V. Yampolskaya and D. H. Walker (1993). “Pathology of inhalational anthrax in 42 cases from the Sverdlovsk outbreak of 1979.” Proc Natl Acad Sci U S A 90(6): 2291–2294.

Affairs, U. N. O. f. D. (2016). “Biological Weapons - The Biological Weapons Convention.” from https://www.un.org/disarmament/wmd/bio/.

Alibek, K. and S. Handelman (1999). “Biohazard: The chilling true story of the largest covert biological weapons in the world-Told from the inside by the man who ran it.” Dell publishing 4: 6.1.

Altschul, S. F., W. Gish, W. Miller, E. W. Myers and D. J. Lipman (1990). “Basic local alignment search tool.” J. Mol. Biol. 215: 403–410.

Archie, J. W. (1989). “Homoplasy excess ratios: new indices for measuring levels of homoplasy in phylogenetic systematics and a critique of the consistency index.” Systematic Biology 38(3): 253–269.

Archie, J. W. (1996). Measures of homoplasy. Homoplasy: The recurrence of similarity in evolution. L. H. M. J. Sanderson. San Diego, Academic Press: 153–188.

Bankevich, A., S. Nurk, D. Antipov, A. A. Gurevich, M. Dvorkin, A. S. Kulikov, V. M. Lesin, S. I. Nikolenko, S. Pham, A. D. Prjibelski, A. V. Pyshkin, A. V. Sirotkin, N. Vyahhi, G. Tesler, M. A. Alekseyev and P. A. Pevzner (2012). “SPAdes: a new genome assembly algorithm and its applications to single-cell sequencing.” J Comput Biol 19(5): 455–477.

Bao, E., T. Jiang and T. Girke (2014). “AlignGraph: algorithm for secondary de novo genome assembly guided by closely related references.” Bioinformatics 30(12): i319–i328.

Benson, D. A., I. Karsch-Mizrachi, K. Clark, D. J. Lipman, J. Ostell and E. W. Sayers (2012). “GenBank.” Nucleic Acids Res 40(Database issue): D48–53.

Bolger, A. M., M. Lohse and B. Usadel (2014). “Trimmomatic: a flexible trimmer for Illumina sequence data.” Bioinformatics 30(15): 2114–2120.

Delcher, A. L., S. L. Salzberg and A. M. Phillippy (2003). “Using MUMmer to identify similar regions in large sequence sets.” Curr Protoc Bioinformatics Chapter 10: Unit 10 13.

DePristo, M. A., E. Banks, R. Poplin, K. V. Garimella, J. R. Maguire, C. Hartl, A. A. Philippakis, G. del Angel, M. A. Rivas, M. Hanna, A. McKenna, T. J. Fennell, A. M. Kernytsky, A. Y. Sivachenko, K. Cibulskis, S. B. Gabriel, D. Altshuler and M. J. Daly (2011). “A framework for variation discovery and genotyping using next-generation DNA sequencing data.” Nature genetics 43(5): 491–498.

Devault, A. M., G. B. Golding, N. Waglechner, J. M. Enk, M. Kuch, J. H. Tien, M. Shi, D. N. Fisman, A. N. Dhody, S. Forrest, K. I. Bos, D. J. D. Earn, E. C. Holmes and H. N. Poinar (2014). “Second-Pandemic Strain of Vibrio cholerae from the Philadelphia Cholera Outbreak of 1849.” New England Journal of Medicine 370(4): 334–340.

Eppinger, M., T. Pearson, S. S. Koenig, O. Pearson, N. Hicks, S. Agrawal, F. Sanjar, K. Galens, S. Daugherty, J. Crabtree, R. S. Hendriksen, L. B. Price, B. P. Upadhyay, G. Shakya, C. M. Fraser, J. Ravel and P. S. Keim (2014). “Genomic epidemiology of the Haitian cholera outbreak: a single introduction followed by rapid, extensive, and continued spread characterized the onset of the epidemic.” MBio 5(6): e01721.

Felsenstein, J. (1985). “Confidence limits on phylogenies: An approach using the bootstrap.” Evolution 39: 783–791.

Hendriksen, R. S., L. B. Price, J. M. Schupp, J. D. Gillece, R. S. Kaas, D. M. Engelthaler, V. Bortolaia, T. Pearson, A. E. Waters, B. P. Upadhyay, S. D. Shrestha, S. Adhikari, G. Shakya, P. S. Keim and F. M. Aarestrup (2011). “Population genetics of Vibrio cholerae from Nepal in 2010: evidence on the origin of the Haitian outbreak.” MBio 2(4): e00157–00111.

Hyatt, D., G. L. Chen, P. F. Locascio, M. L. Land, F. W. Larimer and L. J. Hauser (2010). “Prodigal: prokaryotic gene recognition and translation initiation site identification.” BMC bioinformatics 11: 119.

Jackson, P. J., M. E. Hugh-Jones, D. M. Adair, G. Green, K. K. Hill, C. R. Kuske, L. M. Grinberg, F. A. Abramova and P. Keim (1998). “PCR analysis of tissue samples from the 1979 Sverdlovsk anthrax victims: the presence of multiple Bacillus anthracis strains in different victims.” Proc Natl Acad Sci U S A 95(3): 1224–1229.

Jernigan, D. B., P. L. Raghunathan, B. P. Bell, R. Brechner, E. A. Bresnitz, J. C. Butler, M. Cetron, M. Cohen, T. Doyle and M. Fischer (2002). “Investigation of bioterrorism-related anthrax, United States, 2001: epidemiologic findings.(Bioterrorism-Related Anthrax).” Emerg Infect Dis 8(10): 1019–1029.

Keim, P., R. Grunow, R. Vipond, G. Grass, A. Hoffmaster, D. N. Birdsell, S. R. Klee, S. Pullan, M. Antwerpen, B. N. Bayer, J. Latham, K. Wiggins, C. Hepp, T. Pearson, T. Brooks, J. Sahl and D.M. Wagner (2015). “Whole Genome Analysis of Injectional Anthrax Identifies Two Disease Clusters Spanning More Than 13 Years.” EBioMedicine 2(11): 1613–1618.

Keim, P., L. B. Price, A. M. Klevytska, K. L. Smith, J. M. Schupp, R. Okinaka, P. J. Jackson and M. E. Hugh-Jones (2000). “Multiple-locus variable-number tandem repeat analysis reveals genetic relationships within Bacillus anthracis.” J Bacteriol 182(10): 2928–2936.

Keim, P., T. Pearson, B. Budowle, M. Wilson, D.M. Wagner (2011). Microbial Forensic Investigations in the Context of Bacterial Population Genetics. Microbial Forensics, 2nd Edition.

S. E. S. B. Budowle, R.G. Breeze, P.S. Keim, S.A. Morse, Elsevier: 545–580.

Keim, P., M. N. Van Ert, T. Pearson, A. J. Vogler, L. Y. Huynh and D. M. Wagner (2004). “Anthrax molecular epidemiology and forensics: using the appropriate marker for different evolutionary scales.” Infect Genet Evol 4(3): 205–213.

Kenefic, L. J., T. Pearson, R. T. Okinaka, J. M. Schupp, D. M. Wagner, A. R. Hoffmaster, C. B. Trim, W. K. Chung, J. A. Beaudry, L. Jiang, P. Gajer, J. T. Foster, J. I. Mead, J. Ravel and P. Keim (2009). “Pre-Columbian origins for North American anthrax.” PLoS One 4(3): e4813.

Kent, W. J. (2002). “BLAT--the BLAST-like alignment tool.” Genome Res 12(4): 656–664.

Khmaladze, E., D. N. Birdsell, A. A. Naumann, C. B. Hochhalter, M. L. Seymour, R. Nottingham, S. M. Beckstrom-Sternberg, J. Beckstrom-Sternberg, M. P. Nikolich, G. Chanturia, E. Zhgenti, M. Zakalashvili, L. Malania, G. Babuadze, N. Tsertsvadze, N. Abazashvili, M. Kekelidze, S. Tsanava, P. Imnadze, H. H. Ganz, W. M. Getz, O. Pearson, P. Gajer, M. Eppinger, J. Ravel, D. M. Wagner, R. T. Okinaka, J. M. Schupp, P. Keim and T. Pearson (2014). “Phylogeography of Bacillus anthracis in the country of Georgia shows evidence of population structuring and is dissimilar to other regional genotypes.” Plos One 9(7): e102651.

Kumar, S., G. Stecher and K. Tamura (2016). “MEGA7: Molecular Evolutionary Genetics Analysis version 7.0 for bigger datasets.” Molecular Biology and Evolution.

Leitenberg, M., R. A. Zilinskas and J. H. Kuhn (2012). The Soviet biological weapons program: a history, Harvard University Press.

Li, H. (2013). “Aligning sequence reads, clone sequences and assembly contigs with BWA-MEM.” arXiv.org(arXiv:1303.3997 [q-bio.GN]).

Marston, C. K., C. A. Allen, J. Beaudry, E. P. Price, S. R. Wolken, T. Pearson, P. Keim and A. R. Hoffmaster (2011). “Molecular epidemiology of anthrax cases associated with recreational use of animal hides and yarn in the United States.” Plos One 6(12): e28274.

McKenna, A., M. Hanna, E. Banks, A. Sivachenko, K. Cibulskis, A. Kernytsky, K. Garimella, D. Altshuler, S. Gabriel, M. Daly and M. A. DePristo (2010). “The Genome Analysis Toolkit: a MapReduce framework for analyzing next-generation DNA sequencing data.” Genome research 20(9): 1297–1303.

Meselson, M., J. Guillemin, M. Hugh-Jones, A. Langmuir, I. Popova, A. Shelokov and O. Yampolskaya (1994). “The Sverdlovsk anthrax outbreak of 1979.” Science 266(5188): 1202–1208.

Mock, M. and A. Fouet (2001). Anthrax.” Annu Rev Microbiol 55: 647-–671.

Okinaka, R. T., M. Henrie, K. K. Hill, K. S. Lowery, M. Van Ert, T. Pearson, J. Schupp, L. Kenefic, J. Beaudry, S. A. Hofstadler, P. J. Jackson and P. Keim (2008). “Single nucleotide polymorphism typing of Bacillus anthracis from Sverdlovsk tissue.” Emerg Infect Dis 14(4): 653–656.

Pearson, T., J. D. Busch, J. Ravel, T. D. Read, S. D. Rhoton, J. M. U'Ren, T. S. Simonson, S. M. Kachur, R. R. Leadem, M. L. Cardon, M. N. Van Ert, L. Y. Huynh, C. M. Fraser and P. Keim (2004). “Phylogenetic discovery bias in Bacillus anthracis using single-nucleotide polymorphisms from whole-genome sequencing.” Proc Natl Acad Sci U S A 101(37): 13536–13541.

Pearson, T., R. T. Okinaka, J. T. Foster and P. Keim (2009). “Phylogenetic understanding of clonal populations in an era of whole genome sequencing.” Infect Genet Evol 9(5): 1010–1019.

Pomerantsev, A. P., N. A. Staritsin, V. Mockov Yu and L. I. Marinin (1997). “Expression of cereolysine AB genes in Bacillus anthracis vaccine strain ensures protection against experimental hemolytic anthrax infection.” Vaccine 15(17-18): 1846–1850.

Price, E. P., M. L. Seymour, D. S. Sarovich, J. Latham, S. R. Wolken, J. Mason, G. Vincent, K. P. Drees, S. M. Beckstrom-Sternberg, A. M. Phillippy, S. Koren, R. T. Okinaka, W. K. Chung, J. M. Schupp, D. M. Wagner, R. Vipond, J. T. Foster, N. H. Bergman, J. Burans, T. Pearson, T. Brooks and P. Keim (2012). “Molecular epidemiologic investigation of an anthrax outbreak among heroin users, Europe.” Emerg Infect Dis 18(8): 1307–1313.

Price, L. B., M. Hugh-Jones, P. J. Jackson and P. Keim (1999). “Genetic diversity in the protective antigen gene of Bacillus anthracis.” J Bacteriol 181(8): 2358–2362.

Price, L. B., A. Vogler, T. Pearson, J. D. Busch, J. M. Schupp and P. Keim (2003). “In vitro selection and characterization of Bacillus anthracis mutants with high-level resistance to ciprofloxacin.” Antimicrob Agents Chemother 47(7): 2362–2365.

Pullan, S. T., T. R. Pearson, J. Latham, J. Mason, B. Atkinson, N. J. Silman, C. K. Marston, J. W. Sahl, D. Birdsell, A. R. Hoffmaster, P. Keim and R. Vipond (2015). “Whole-genome sequencing investigation of animal-skin-drum-associated UK anthrax cases reveals evidence of mixed populations and relatedness to a US case.” Microbial Genomics 1(5).

Rambaut, A., T. T. Lam, L. Max Carvalho and O. G. Pybus (2016). “Exploring the temporal structure of heterochronous sequences using TempEst (formerly Path-O-Gen).” Virus Evolution 2(1).

Ross, C. L., K. S. Thomason and T. M. Koehler (2009). “An extracytoplasmic function sigma factor controls beta-lactamase gene expression in Bacillus anthracis and other Bacillus cereus group species.” J Bacteriol 191(21): 6683–6693.

Sahl, J. W., S. M. Beckstrom-Sternberg, J. Babic-Sternberg, J. D. Gillece, C. M. Hepp, R. K. Auerbach, W. Tembe, D. M. Wagner, P. S. Keim and T. Pearson (2015). “The In Silico Genotyper (ISG): an open-source pipeline to rapidly identify and annotate nucleotide variants for comparative genomics applications.” bioRxiv.

Sahl, J. W., D. Lemmer, J. Travis, J. Schupp, J. Gillece, M. Aziz, E. Driebe, K. Drees, N. D. Hicks, C. Williamson, C. Hepp, D. E. Smith, C. Roe, D. M. Engelthaler, D. M. Wagner and P. Keim (2016). “The Northern Arizona SNP Pipeline (NASP): accurate, flexible, and rapid identification of SNPs in WGS datasets.” bioRxiv.

Sahl, J. W., J. M. Schupp, D. A. Rasko, R. E. Colman, J. T. Foster and P. Keim (2015).“Phylogenetically typing bacterial strains from partial SNP genotypes observed from direct sequencing of clinical specimen metagenomic data.” Genome Med 7(1): 52.

Stepanov, A. V., L. I. Marinin, A. P. Pomerantsev and N. A. Staritsin (1996). “Development of novel vaccines against anthrax in man.” J Biotechnol 44(1-3): 155–160.

Takahashi, H., P. Keim, A. F. Kaufmann, C. Keys, K. L. Smith, K. Taniguchi, S. Inouye and T. Kurata (2004). “Bacillus anthracis incident, Kameido, Tokyo, 1993.” Emerg Infect Dis 10(1): 117–120.

Tigertt, W. D. (1980). “Anthrax. William Smith Greenfield, M.D., F.R.C.P., Professor Superintendent, the Brown Animal Sanatory Institution (1878-81). Concerning the priority due to him for the production of the first vaccine against anthrax.” J Hyg (Lond) 85(3): 415–420.

Van Ert, M. N., W. R. Easterday, L. Y. Huynh, R. T. Okinaka, M. E. Hugh-Jones, J. Ravel, S. R. Zanecki, T. Pearson, T. S. Simonson, J. M. U'Ren, S. M. Kachur, R. R. Leadem-Dougherty, S. D. Rhoton, G. Zinser, J. Farlow, P. R. Coker, K. L. Smith, B. Wang, L. J. Kenefic, C. M. Fraser-Liggett, D. M. Wagner and P. Keim (2007). “Global genetic population structure of Bacillus anthracis.” Plos One 2(5): e461.

Vergnaud, G., G. Girault, S. Thierry, C. Pourcel, N. Madani and Y. Blouin (2016). “Comparison of French and Worldwide Bacillus anthracis Strains Favors a Recent, Post-Columbian Origin of the Predominant North-American Clade.” PLoS One 11(2): e0146216.

Vogler, A. J., J. D. Busch, S. Percy-Fine, C. Tipton-Hunton, K. L. Smith and P. Keim (2002). “Molecular analysis of rifampin resistance in Bacillus anthracis and Bacillus cereus.” Antimicrob Agents Chemother 46(2): 511–513.

Walker, B. J., T. Abeel, T. Shea, M. Priest, A. Abouelliel, S. Sakthikumar, C. A. Cuomo, Q. Zeng, J. Wortman, S. K. Young and A. M. Earl (2014). “Pilon: an integrated tool for comprehensive microbial variant detection and genome assembly improvement.” PLoS One 9(11): e112963.

Wilgenbusch, J. C. and D. Swofford (2003). “Inferring evolutionary trees with PAUP*.” Curr Protoc Bioinformatics Chapter 6: Unit 6 4.

Zwick, M. E., S. J. Joseph, X. Didelot, P. E. Chen, K. A. Bishop-Lilly, A. C. Stewart, K. Willner, N. Nolan, S. Lentz, M. K. Thomason, S. Sozhamannan, A. J. Mateczun, L. Du and T. D. Read (2012). “Genomic characterization of the Bacillus cereus sensu lato species: backdrop to the evolution of Bacillus anthracis.” Genome research 22(8): 1512–1524.

